# Antimuscarinic drugs exert β-arrestin-biased agonism at the muscarinic acetylcholine type 1 receptor

**DOI:** 10.1101/2025.04.16.649213

**Authors:** Shayan Amiri, Mohamad-Reza Aghanoori, Darrell R. Smith, T. M. Zaved Waise, Ying Lao, Asuka Inoue, René P. Zahedi, Henry A. Dunn, Paul Fernyhough

**Author notes:** Corresponding author: Dr. Paul Fernyhough, Department of Pharmacology & Therapeutics, Rady Faculty of Health Sciences, University of Manitoba. Current address of Dr. Aghanoori: Department of Molecular Medicine, National Institute of Genetic Engineering and Biotechnology (NIGEB), Tehran, Iran. Current address of Dr. Waise: Department of Zoology, 3112 - 6270 University Blvd., University of British Columbia, Vancouver, BC, V6T 1Z4, Canada.

## Abstract

Previous studies indicate that both pirenzepine (PZ), a selective orthosteric muscarinic acetylcholine type 1 receptor (M_1_R) antagonist, and muscarinic toxin 7 (MT7), a negative M_1_R allosteric modulator (NAM), act via M_1_R to promote neuritogenesis in cultured adult rodent primary dorsal root ganglia (DRG) sensory neurons, in part, through β-arrestin-dependent activation of extracellular signal-regulated protein kinase 1/2 (ERK1/2). Furthermore, these antagonists reverse nerve degeneration in a variety of rodent models of peripheral neuropathy through multiple complementary pathways. To understand the therapeutic effects and mechanism of M_1_R antagonist-induced ERK1/2 phosphorylation, we tested the hypothesis that PZ and MT7 possess β−arrestin-biased agonism at M_1_R to drive activation of ERK and enhance neurite outgrowth. Treatment for up to 30 min with PZ and MT7 dose-dependently recruited β-arrestin2 to M_1_R (analyzed using nano-BRET) and increased ERK phosphorylation in both HEK293 cells and DRG neurons. DRG neurons of different sub-types express M_1_R, and ERK activation by MT7 was only observed in M_1_R-positive neurons. These novel pharmacological effects occurred in the absence of activation of G protein signaling or receptor internalization. PZ phosphorylated M_1_R at six specific serine/threonine residues (T230, S251, T254, S321, T354, S356) of intracellular loop 3 (ICL3) and deletion mutation of these sites suppressed PZ and MT7 induction of β-arrestin binding to M_1_R and inhibited ERK activation. With regard to PZ signaling, alanine substitution at S251 and T254 was sufficient to impede β-arrestin binding and ERK activation. β-arrestin-biased activity of PZ and MT7 involved the mobilization of casein kinase 2 (CK2) and this occurred in the absence of Gαq or G protein receptor kinase (GRK) activity. Pharmacological or siRNA-based inhibition of CK2 blocked PZ-induction of β-arrestin association, ERK activation and neurite outgrowth in DRG neurons. In conclusion, PZ/MT7 activated M_1_R toward the β-arrestin signaling pathway in both HEK293 cells and DRG neurons to augment ERK activation and neurite outgrowth via engagement of CK2.

**One-sentence summary:** Antimuscarinic drugs act as β-arrestin-biased agonists via casein kinase 2 activation to promote ERK1/2 phosphorylation and neurite outgrowth

Graphical abstract.
Schematic presentation of the effect of muscarinic ligands at M_1_R associated signaling pathway.
**(A)** Muscarine/carbachol acts as a balanced ligand by engaging both Gα_q_ and β-arrestin signaling pathways and treatment with pirenzepine/MT7 blocks these effects. **(B)** Pirenzepine/MT7 acts as a β-arrestin biased ligand by 1) phosphorylating of ICL3 region of M_1_R via CK2 (but not GRKs), 2) no activation of G protein signaling, 3) recruitment of β-arrestin 2 and 4) ERK1/2 activation leading to neurite outgrowth in DRG sensory neurons. This figure was generated by BioRender under license number EK285MUOPQ.

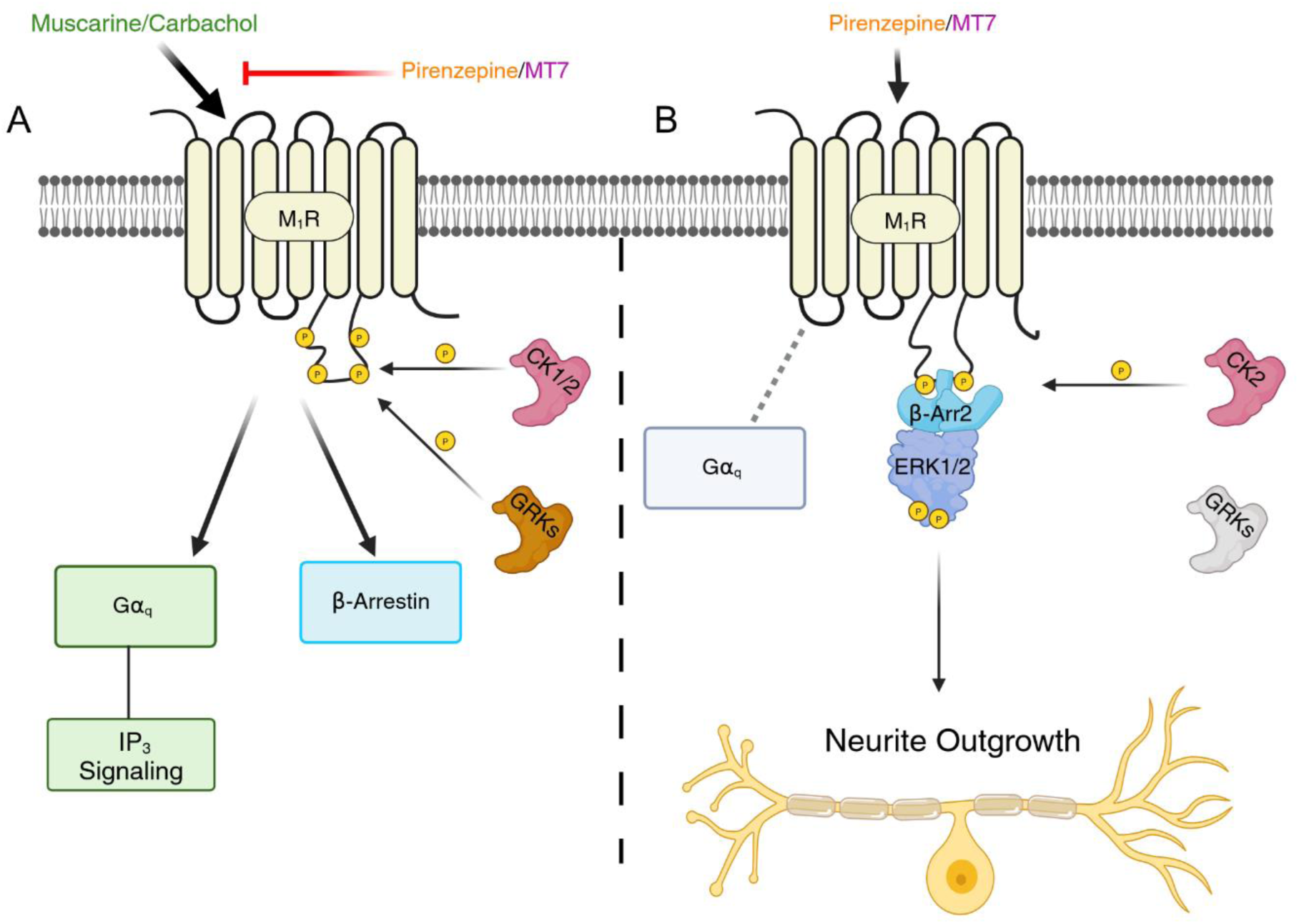

## INTRODUCTION

G protein–coupled receptors (GPCRs) are dynamic transmembrane proteins that activate several signaling pathways via engagement of numerous intracellular molecules, including G proteins, G protein-coupled receptor kinases (GRKs), and β-arrestins (*1*). Recruitment and activation of the different heterotrimeric G proteins and β-arrestins are ligand-dependent and cell-type-dependent processes (*2*). Ligand binding to GPCRs triggers phosphorylation events at intracellular loops (ICLs) and the carboxyl terminus of the receptors that mobilizes binding of β-arrestins. These phosphorylation events are mostly controlled by kinases such as GRKs (*3*) and casein kinases (*4–6*). It is now clear that unlike balanced agonists and antagonists, biased ligands can specifically target a particular receptor-linked signaling pathway or subset of pathways (*7, 8*).

The muscarinic acetylcholine type 1 receptor (M_1_R) is a class A GPCR and widely expressed in the central nervous system (CNS) (*9, 10*) and the peripheral nervous system (PNS) (*11, 12*). In response to the endogenous agonist, acetylcholine, or a balanced synthetic agonist, such as carbachol, M_1_R activates several signaling pathways via recruitment of Gα_q_/_11_ to increase inositol (1,4,5) trisphosphate (IP_3_) and intracellular calcium levels, and β-arrestins contributing to activation of extracellular signal-regulated protein kinases (ERKs) (*13–16*). M_1_R plays an important role in the regulation of neuronal function and its ablation causes absence of phosphoinositide hydrolysis in cortical and hippocampal neurons in response to muscarinic agonists (*17, 18*). M_1_R is the major muscarinic receptor subtype that mediates ERK activation in neurons, and M_1_R knockout (KO) mice show no ERK activation in hippocampal and cortical neurons after receiving carbachol and oxotremorine, respectively (*19, 20*). M_1_R selective agonists promote cellular excitability, synaptic transmission, and hippocampal long-term potentiation in mice, and this is suppressed in M_1_R KO mice (*21*). A study using chemogenetics and a phospho-deficient M_1_R mouse model revealed that regulation of learning, memory, and anxiety-like behaviors depends on M_1_R phosphorylation and relevant signaling pathways (such as β-arrestin recruitment and ERK activation), while Gα_q_-related signaling was linked to M_1_R-dependent adverse effects such as seizures (*22*).

We previously reported that genetic or pharmacological blockade of M_1_R exhibited protection against different types of peripheral neuropathy in rodents, including sensory and motor nerve dysfunction (*23*). Treatment with pirenzepine (PZ), a selective orthosteric antagonist for M_1_R, and muscarine toxin 7 (MT7), a negative allosteric modulator (NAM) for M_1_R, enhanced mitochondrial function and neurite outgrowth in adult rodent dorsal root ganglia (DRG) sensory neurons (*16, 23*). Further, we showed that ERK activation and β-arrestins contribute to the neuritogenic effect of PZ and MT7 (*24*). However, the molecular mechanisms through which these prototypical antagonists activate M_1_R-mediated ERK1/2 remain unknown. Interestingly, prototypical β-adrenergic receptor antagonists, such as the prominent β-blocker carvedilol, have also demonstrated β-arrestin-biased agonism leading to the induction of ERK activation (*25, 26*). Studies with the β2-adrenergic receptor have assigned a barcode related to the distinct phosphorylation pattern controlled by GRKs that determines β-arrestin-dependent signaling (*26*). Thus, in the current study, we tested the hypothesis that PZ and MT7 may possess β-arrestin-biased agonism at the M_1_R.

To address this hypothesis, we used an orthosteric antagonist (PZ) and a NAM (MT7) to study signaling pathways and molecular mechanisms associated with M_1_R, including G protein signaling, β-arrestin recruitment, ERK activation, receptor phosphorylation and internalization. We found that M_1_R antagonists phosphorylated M_1_R at specific residues at intracellular loop 3 (ICL3), recruited β-arrestin2, and activated ERK in both HEK293 and primary adult DRG sensory neurons without activating G protein signaling or receptor internalization. ERK activation depended on the phosphorylation of specific residues at M_1_R. Unlike Gα_q_ or GRKs, we found that β-arrestins and casein kinase 2 (CK2) mediate the effects of antimuscarinic drugs on ERK activation. Finally, inhibition of CK2 blocked PZ-induced ERK activation and neurite outgrowth in adult DRG sensory neurons.

## RESULTS

### Pirenzepine (PZ) and MT7 recruited β-arrestin2 to M_1_R without activation of Gα_q_ protein signaling

To explore whether MT7 and PZ mobilize β-arrestin signaling in the absence of G protein signaling we first showed the classical antagonism activity of PZ and MT7 at M_1_R by measuring inositol monophosphate (IP1) production. As seen in Fig. 1A&B, PZ and MT7 blocked muscarine-induced IP1 production in a dose-dependent manner in HEK293 cells and cultured adult rat DRG sensory neurons (Fig. 3B&C). Next, we examined the induction of β-arrestin2 recruitment to M_1_R using a Bioluminescence Resonance Energy Transfer (BRET) assay. The rationale for selecting β-arrestin2 over β-arrestin1 was that M_1_R has more preference for β-arrestin2 (*27, 28*). BRET measurements showed that treatment with carbachol (1 mM) (Fig. S1) or muscarine (1 mM) rapidly increased β-arrestin2 recruitment to M_1_R (Fig. 1C&D). Subsequent treatment over a 5 min period with the PZ (1 µM), or MT7 (100 nM) inhibited the muscarine-induced interaction between M_1_R and β-arrestin2 (Fig. 1C&D); as expected and previously described (*15*). Further, we showed that PZ and MT7 treatments (but not muscarine) failed to increase IP1 in HEK293 cells (Fig. 2A). Interestingly, unlike muscarine, PZ and MT7 treatments did not change the cAMP response to forskolin in HEK293 cells (Fig. 2B). Since the maximum BRET response to M_1_R agonists were observed after 5 min of incubation, a dose-response curve was performed at this time point. M_1_R activation induced by carbachol over a 5 min period induced a dose-dependent recruitment of β-arrestin2 in HEK293 cells (Fig. 2C) and DRG sensory neurons (Fig. 3D). Unexpectedly, however, over a longer-term treatment period of 30 min PZ recruited β-arrestin2 to M_1_R in a concentration-dependent manner in HEK293 cells (Fig. 2D) and DRG neurons (Fig. 3E). Similarly, over a 30 min period MT7 induced recruitment of β-arrestin2 to M_1_R in a dose-dependent manner in HEK293 cells (Fig. 2E) and DRG sensory neurons (Fig. 3F).

**Fig. 1.**
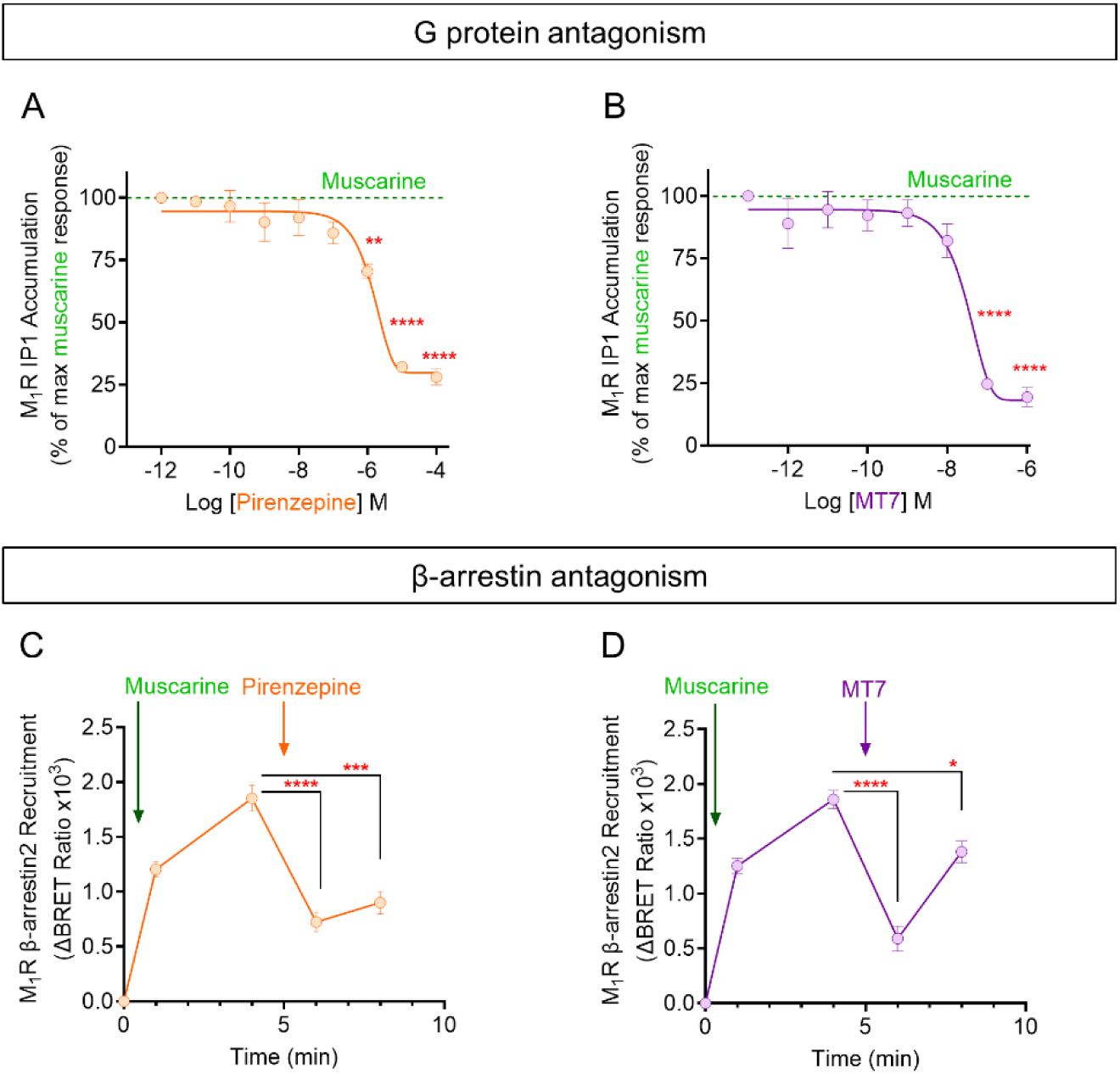
Antagonism activity of pirenzepine and MT7 against muscarine-induced signaling at M_1_R. **(A)** Inhibitory effect of different doses of pirenzepine and **(B)** MT7 on muscarine (1 mM)-induced IP1 production in transiently hM_1_R-transfected HEK293 cells. Data show mean ± SEM (1-way ANOVA with Dunnett’s post-hoc test). Data normalized to baseline with multiple comparisons to the lowest concentration of drug; ****P<0.0001, **P<0.01, n=3. **(C&D)** Ligands were added at the times indicated. Muscarine (1 mM) induced βArr2-Halo recruitment to hM_1_R-Nluc, and acute treatment with **(C)** pirenzepine (1 µM) and **(D)** MT7 (100 nM) blocked the effects of muscarine in transiently co-transfected HEK293 cells. Data show mean ± SEM (1-way ANOVA with Dunnett’s post-hoc test). Antagonist-treated groups were compared to muscarine-treated groups at minute 4; ****P<0.0001, ***P<0.001, and *P<0.05, n=3.

**Fig. 2.**
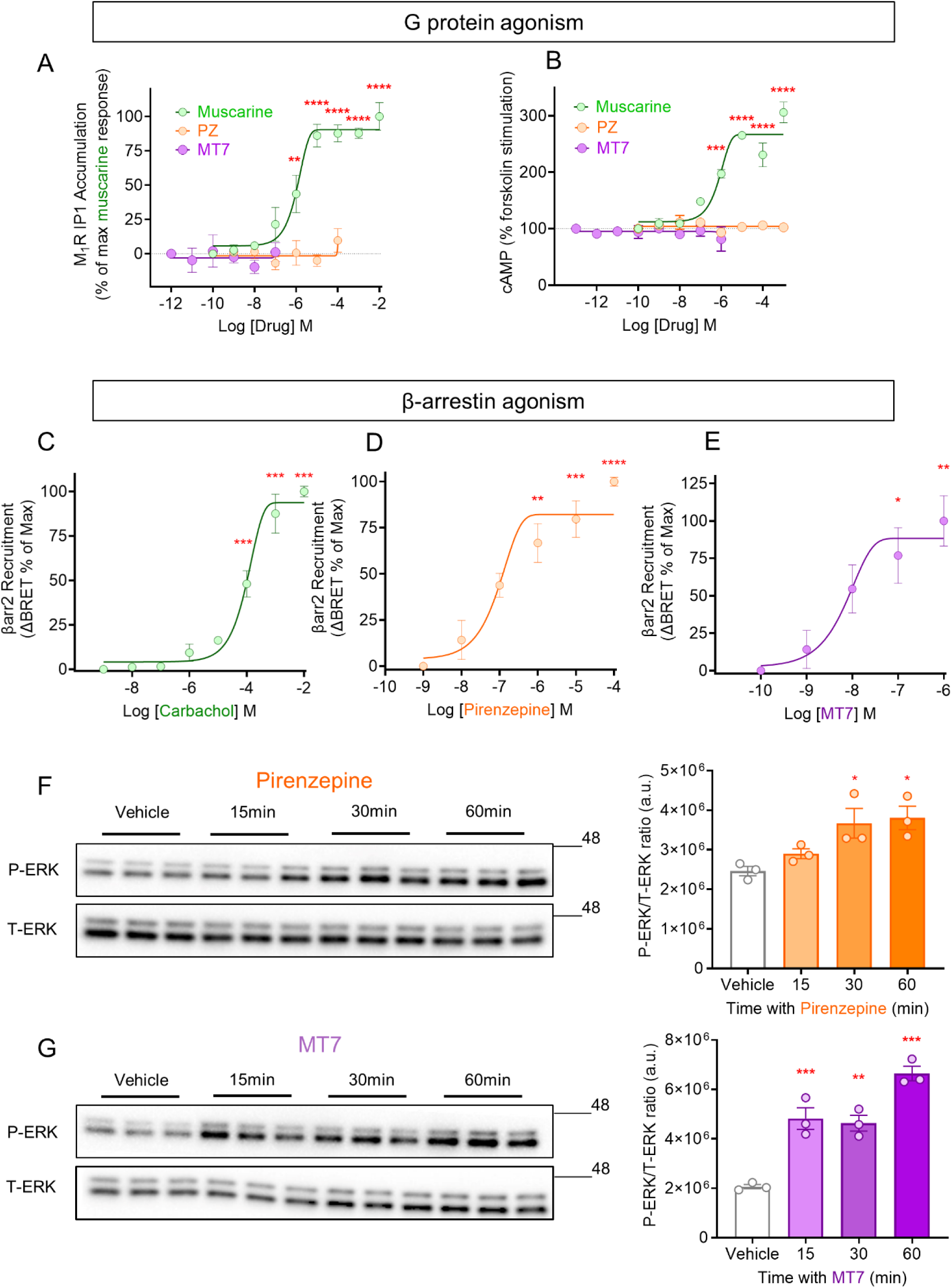
Pirenzepine and MT7 activated β-arrestin (but not G protein) pathway in HEK293 cells. **(A)** shows a dose-response curve for treatment with muscarine, pirenzepine (PZ), and MT7 and levels of IP1 production in hM_1_R-transfected HEK293 cells (at 45 min of treatment). **(B)** shows a dose-response curve for the effects of ligands (muscarine (5 min), PZ (30 min), and MT7 (30 min)) on forskolin (300 nM)-induced cAMP production in hM_1_R-transfected HEK293 cells. Data show mean ± SEM (1-way ANOVA with Dunnett’s post-hoc test). Data normalized to baseline with multiple comparisons to the lowest concentration of drug; ****P<0.001, ***P<0.001, and **P<0.01, n=4. **(C-E)** Recruitment of HaloTag-β-arrestin2 to hM_1_R in transiently co-transfected HEK293 cells following treatments with **(C)** carbachol (5 min), **(D)** pirenzepine (30 min) and **(E)** MT7 (30 min). Data show mean ± SEM (1-way ANOVA with Dunnett’s post-hoc test). Data normalized to baseline with multiple comparisons to the lowest concentration of drug; ****P<0.0001, ***P<0.001, **P<0.01 and *P<0.05, n=3. **(F&G)** Immunoblotting for T-ERK and P-ERK proteins was conducted on hM_1_R**-**transfected HEK293 cells, which were treated with **(F)** pirenzepine (PZ, 1 µM) or **(G)** MT7 (100 nM) at different time-points. Data represent P-ERK/T-ERK ratio (a.u.). Data show mean ± SEM (1-way ANOVA with Dunnett’s post-hoc test). Comparison to vehicle-treated group; ***P<0.001, **P<0.01, and *P<0.05, n=3.

**Fig. 3.**
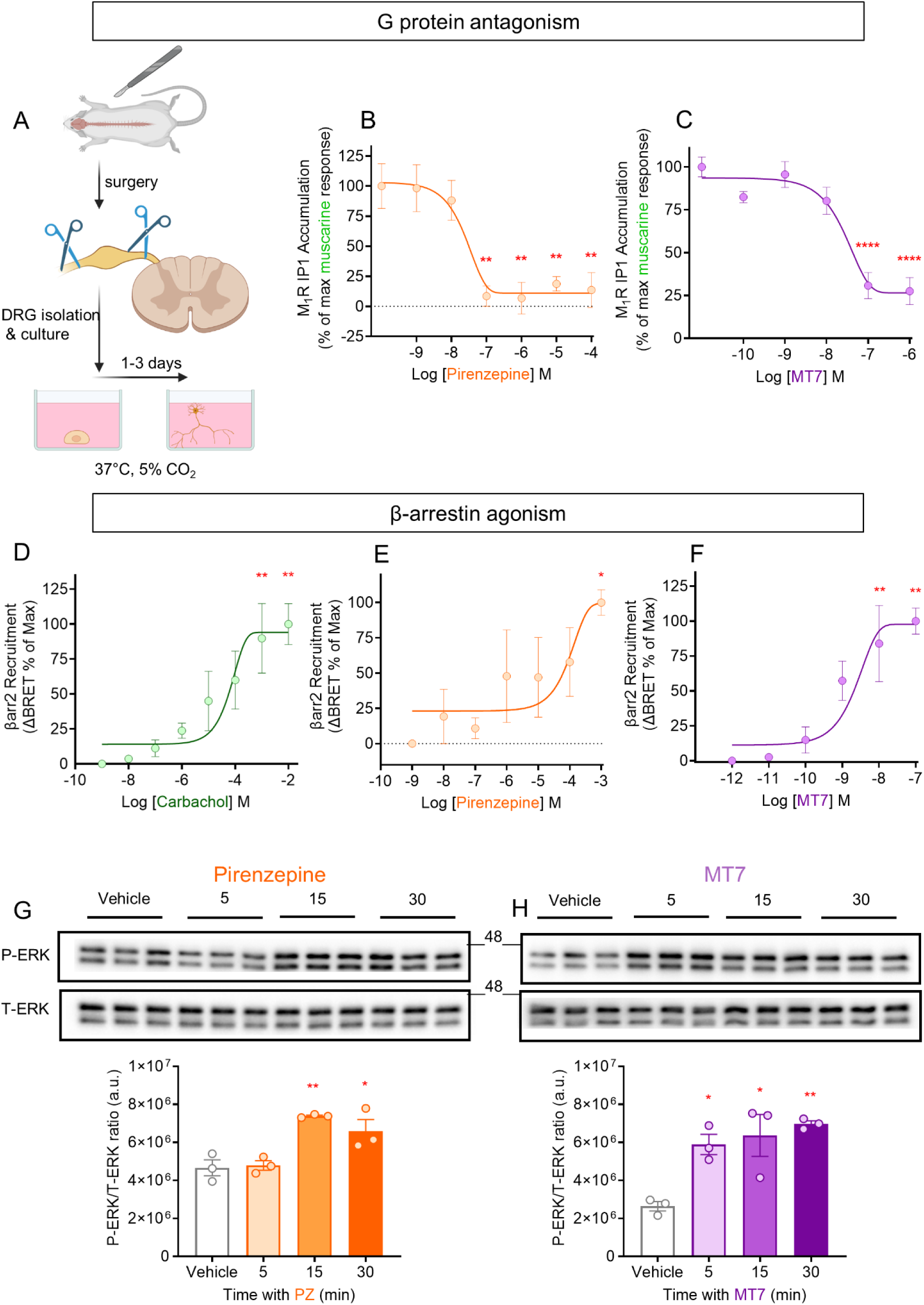
Pirenzepine and MT7 activated β-arrestin (but not G protein) pathway in DRG sensory neurons. **(A)** shows a schematic procedure for adult rat dorsal root ganglion (DRG) neuron isolation and culture. Inhibitory effect of **(B)** different concentrations of pirenzepine and **(C)** MT7 on muscarine (1 mM)-induced IP1 production in DRG sensory neurons. Data show mean ± SEM (1-way ANOVA with Dunnett’s post-hoc test). Data normalized to baseline with multiple comparisons to the lowest concentration of drug; ****P<0.0001, **P<0.01, n=3. **(D-F)** Recruitment of HaloTag-β-arrestin2 to hM_1_R in transiently co-transfected DRG sensory neurons following treatments with **(D)** carbachol (5 min), **(E)** pirenzepine (30 min) and **(F)** MT7 (30 min). Data show mean ± SEM (1-way ANOVA with Dunnett’s post-hoc test). Data normalized to baseline with multiple comparisons to the lowest concentration of drug; **P<0.01, and *P<0.05, n=3. **(G&H)** Immunoblotting for T-ERK and P-ERK proteins was conducted on DRG sensory neurons, which were treated with **(G)** pirenzepine (PZ, 1 µM) or **(H)** MT7 (100 nM) at different time-points. Data represent P-ERK/T-ERK ratio (a.u.). Data show mean ± SEM (1-way ANOVA with Dunnett’s post-hoc test). Comparison to vehicle-treated group **P<0.01, and *P<0.05, n=3.

### PZ and MT7 increased ERK activity in HEK293 cells and DRG neurons

β-arrestins form signaling scaffolds with members of mitogen-activated protein kinases (MAPKs) including ERK (*28, 29*). We previously reported that treating DRG neurons with PZ (1 µM) or MT7 (100 nM) activated ERK via a β-arrestin-dependent pathway and increased neurite outgrowth (*24*). Treating M_1_R-transfected HEK293 cells (Fig. 2F&G) and DRG neurons (Fig. 3G&H) with PZ (1µM) and MT7 (100 nM) increased ERK phosphorylation in a time-dependent manner. To examine the role of M_1_R in the PZ and MT7 stimulation of ERK phosphorylation in DRG neurons, we first stained intact DRG from WT (Fig. 4A, top panel) and M_1_R KO mice (Fig. 4A, bottom panel) with β-tubulin III and MT7-ATTO590 (a functional fluorescently labelled MT7). Since MT7 possesses an extreme subtype specificity toward M_1_R (*30*), we used the MT7-ATTO590 as an M_1_R marker in DRG neurons instead of unreliable commercial antibodies against M_1_R. DRG neurons across a range of perikaryal sizes (and thus sensory sub-types) in WT mice expressed M_1_R (Fig. 4A, top panel) while we confirmed M_1_R KO mice expressed no M_1_R (Fig. 4A, bottom panel). Treating DRG neurons with MT7-ATTO590 (90 min) significantly increased phospho-ERK expression in DRG neurons, but only in those expressing M_1_R (Fig. 4B). The phospho-ERK signal was stronger in the nucleus than cytoplasm of DRG neurons (Fig. 4B).

**Fig. 4.**
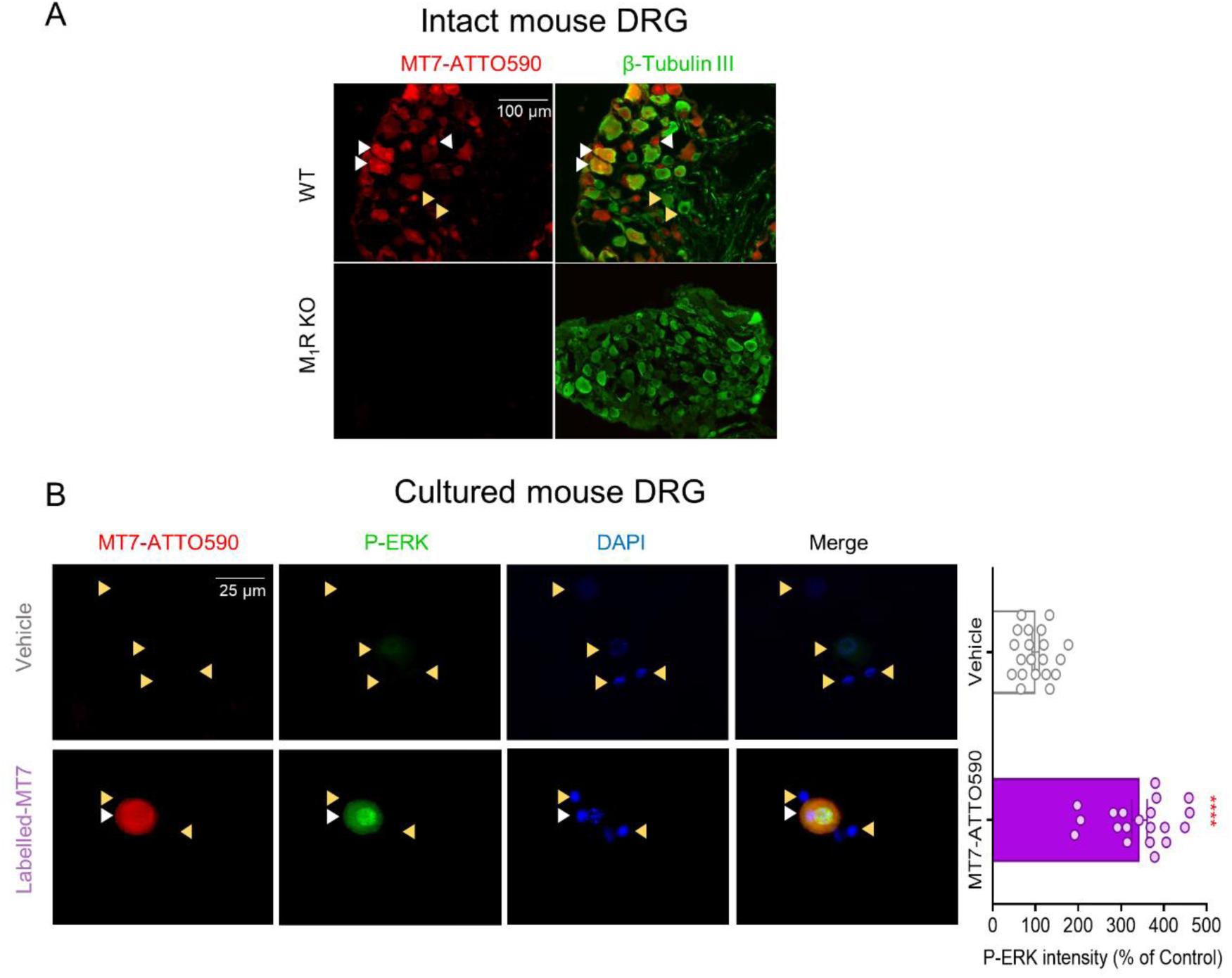
M_1_R expression and single cell analysis of ERK activation in mouse DRG sensory neurons. **(A)** Immunocytochemistry images of adult mouse DRG tissue sections for wild-type (WT) and M_1_R KO mice treated with fluorescently tagged MT7-ATTO590 (100 nM), and labelled with antibody to neuron-specific ß-tubulin III. White arrows indicate M_1_R positive and yellow arrows indicate M_1_R negative neurons. **(B)** Immunocytochemistry images of cultured adult mouse DRG sensory neurons with vehicle/ MT7-ATTO590 treatment (100 nM, 90 min). Neurons were labelled with MT7-ATTO590, phospho-ERK, and DAPI. Values for fluorescence intensity of P-ERK are expressed as the mean ± SEM from 20 neurons and analyzed using *t*-test, ****P<0.0001.

### M_1_R phosphorylation by PZ is necessary for β-arrestin2 biased signaling activity

GPCR function is regulated by phosphorylation (*31*), and β-arrestins bind to phosphorylated GPCRs with high affinity (*32*). To determine whether PZ phosphorylated M_1_R, and to establish the exact sites of phosphorylation, we performed liquid chromatography-tandem mass spectrometry (LC-MS/MS). Results from 2 different proteomics centers (Fig. S2A&B, Table S1, University of Virginia Biomedical Research Facility for LC/MS/MS analysis) and (Fig. S3A-F, Table S2, Manitoba Centre for Proteomics and Systems Biology (MCPSB)) identified three serine (Ser^251^, Ser^321^, Ser^356^) and three threonine (Thr^230^, Thr^254^, Thr^354^) phosphorylation sites in the ICL3 of M_1_R (Fig. 5A). For the MCPSB analysis accurate determination of the phosphorylation degree at individual sites required absolute quantification of both the phosphorylated and non-phosphorylated variants of the corresponding peptides, as previously described (*33*). In this study, we approximated phosphorylation levels by calculating, for each site, the ratio of high-confidence peptide-spectrum matches (PSMs) with site localization probabilities >90% to the total number of PSMs identifying peptides that included the respective amino acid residue. Across four control samples, Ser^251^ and Thr^254^ emerged as the most prominently phosphorylated residues, with 12.8% and 15.5% of PSMs phosphorylated, respectively, while all other sites ranged between 4.4% and 8.8%. Based on these LC-MS/MS results, 2 mutated M_1_R plasmids were generated; 1) deletion of all six phosphorylation sites from ICL3 region of M_1_R (M_1_RΔICL3), and 2) mutation at Ser^251^ and Thr^254^ (M_1_R-S251A, T254A; alanine substitutions) in the ICL3 region. When compared to M_1_R-WT, BRET results showed that M_1_RΔICL3 and M_1_R-S251A, T254A mutations significantly decreased the efficacy (Emax response) of PZ to induce β-arrestin2 recruitment to M_1_R (Fig. 5B). These mutations resulted in the failure of PZ to induce ERK phosphorylation in HEK293 cells (Fig. 6A&B). In comparison to M_1_R-WT, M_1_RΔICL3 (but not M_1_R-S251A, T254A) significantly decreased the efficacy of MT7 to induce M_1_R-β-arrestin2 interaction (Fig. 5C) and ERK activation in HEK293 cells (Fig. 6C&D). Intriguingly, both mutations had no significant effect on the efficacy of muscarine to recruit β-arrestin2 to M_1_R (Fig. 5D). Surprisingly, these mutations significantly increased ERK phosphorylation in response to muscarine, when compared to the M_1_R-WT response (Fig. 6E&F).

**Figure 5.**
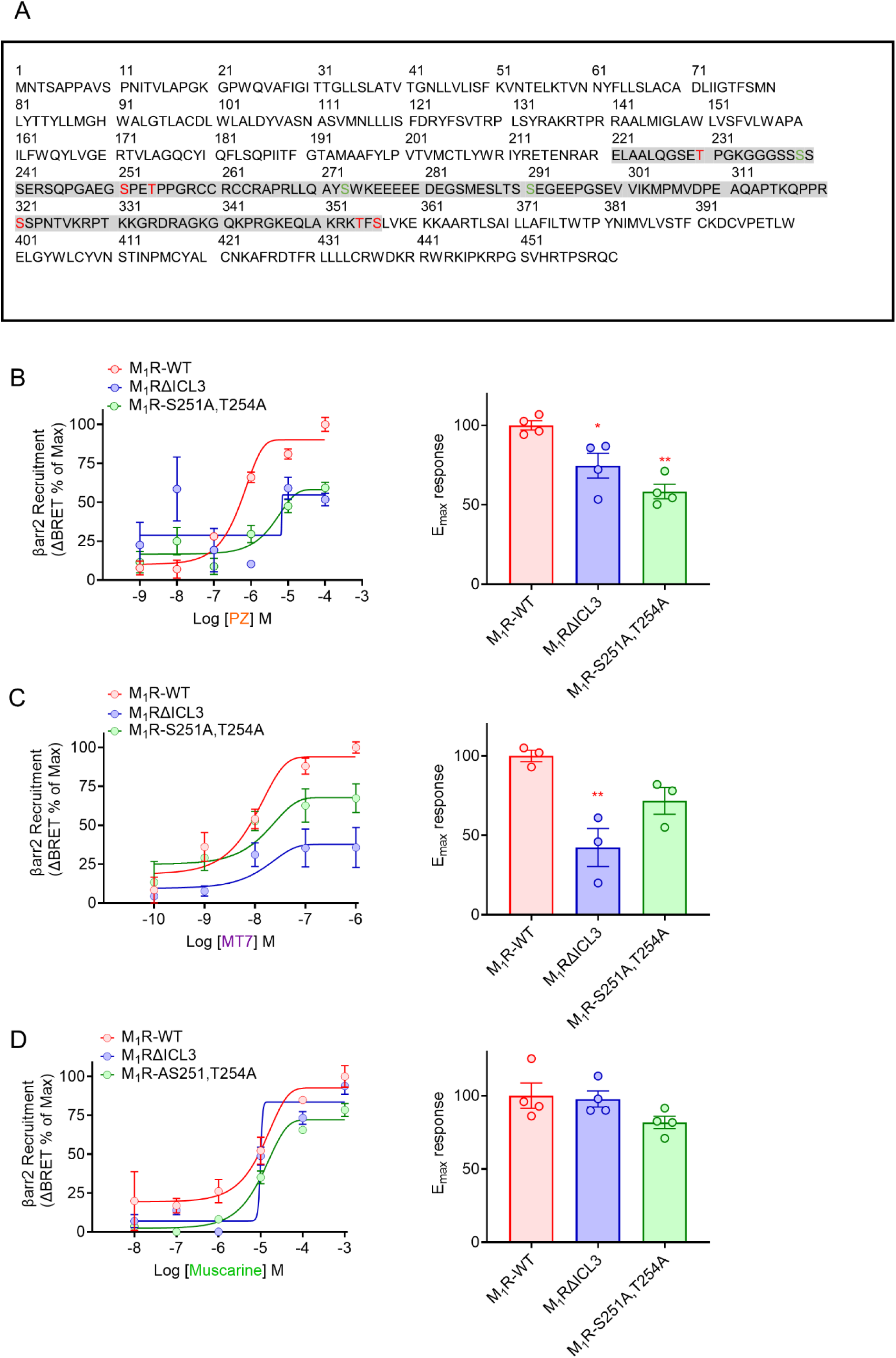
Mass spectrometry for Ser/Thr phosphorylation at hM_1_R and effects of mutagenesis studies on Emax response to pirenzepine. **(A)** M_1_R serine and threonine phosphorylation sites identified by nano-LC-MS/MS with high confidence and with site localization probabilities >0.99. Phosphorylated residues are in the red color within ICL3 region of M_1_R in gray color. Concentration-response results for **(B)** pirenzepine (PZ, 30 min), **(C)** MT7 (30 min) and **(D)** muscarine (5 min) for β-arrestin2 recruitment to M_1_R-WT, M_1_RΔICL3 (mutation of all 6 Phos-sites at M_1_R ICL3 region), and M_1_R-S251A, T254A (mutations at S251 and T254 sites at M_1_R ICL3 region) in HEK293 cells. Emax response was calculated for each drug and data show mean ± SEM (1-way ANOVA with Dunnett’s post-hoc test). Comparisons to M_1_R-WT group; **P<0.01 and *P<0.05, n=4.

**Fig. 6.**
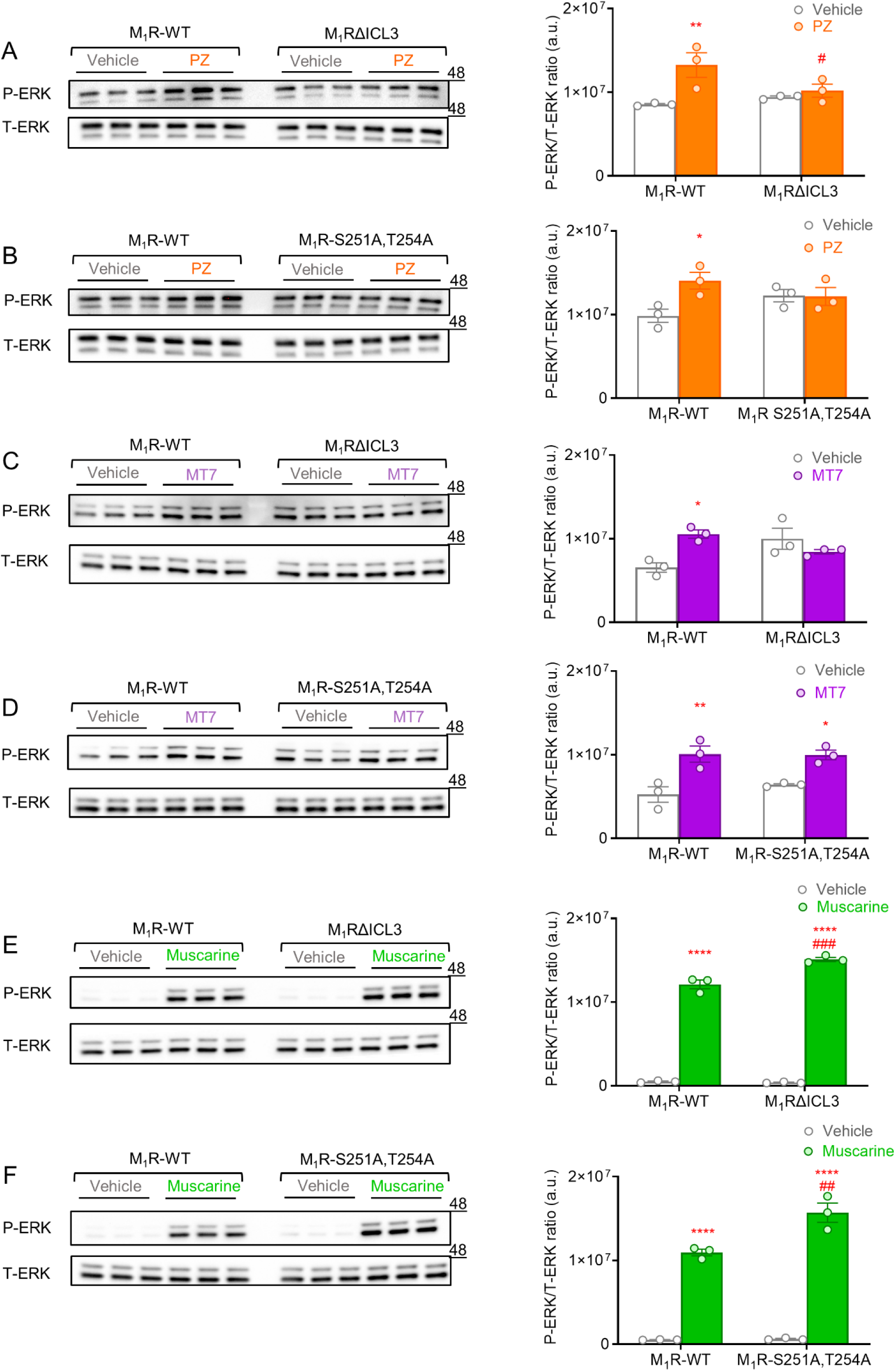
M_1_R mutations altered ERK phosphorylation in response to muscarinic drugs in HEK293 cells. (A, C, and. **E)** HEK293 cells were transfected with M_1_R-WT and M_1_RΔICL3 (mutation of all 6 Phos-sites at M_1_R ICL3 region) and received **(A)** pirenzepine (PZ, 1 µM, 60 min), **(C)** MT7 (100 nM, 60 min), and **E**) muscarine (100 µM, 5 min). **(B, D, and F)** HEK293 cells were transfected with M_1_R-WT and M_1_R-S251A, T254A (mutations at S251 and T254) and received **(B)** pirenzepine (PZ, 1 µM, 60 min), **(D)** MT7 (100 nM, 60 min), and **(F)** muscarine (100 µM, 10 min). Immunoblotting for T-ERK and P-ERK proteins was conducted. Data represent P-ERK/T-ERK ratio (a.u.). Data show mean ± SEM (2-way ANOVA with Tukey’s post-hoc test). Comparison between vehicle-treated groups and drug-treated groups for WT/mutation conditions; ****P<0.0001, **P<0.01, and *P<0.05. Comparison between PZ-treated groups; #P<0.05, comparison between muscarine-treated groups; ###P<0.001 and ##P<0.01, n=3.

### PZ and MT7 increased cell surface expression of M_1_R at HEK293 cells and DRG neurons

It has been known for more than 20 years that β-arrestins bind phosphorylated GPCRs and contribute to the desensitization and internalization of GPCRs (*34*). To assess receptor internalization in response to muscarinic drugs, we used HEK293 cells and DRG sensory neurons transfected with N-terminally HiBiT-tagged M_1_R. Muscarine, as an agonist, dose-dependently increased M_1_R internalization in both HEK293 cells (Fig. 7A) and DRG sensory neurons (Fig. 7D) after 1 h, when compared to vehicle-treated counterparts. However, 1 h of treatment with different doses of PZ increased M_1_R cell surface expression in a concentration-dependent manner in HEK293 cells, when compared to vehicle-treated groups (Fig. 7B). Similar treatment with different PZ concentrations did not significantly reduce/increase M_1_R surface expression in cultured DRG sensory neurons (Fig. 7E). Treating HEK293 cells (Fig. 7C) and DRG sensory neurons (Fig. 7F) with MT7 for 1 h resulted in a dose-dependent increase in cell surface expression of M_1_R. Also, treating HEK293 cells and DRG neurons with PZ (1 µM, Fig. S4A&C) and MT7 (100 nM, Fig. S4B&D) for up to 3 h did not reduce the surface expression of M_1_R.

**Fig. 7.**
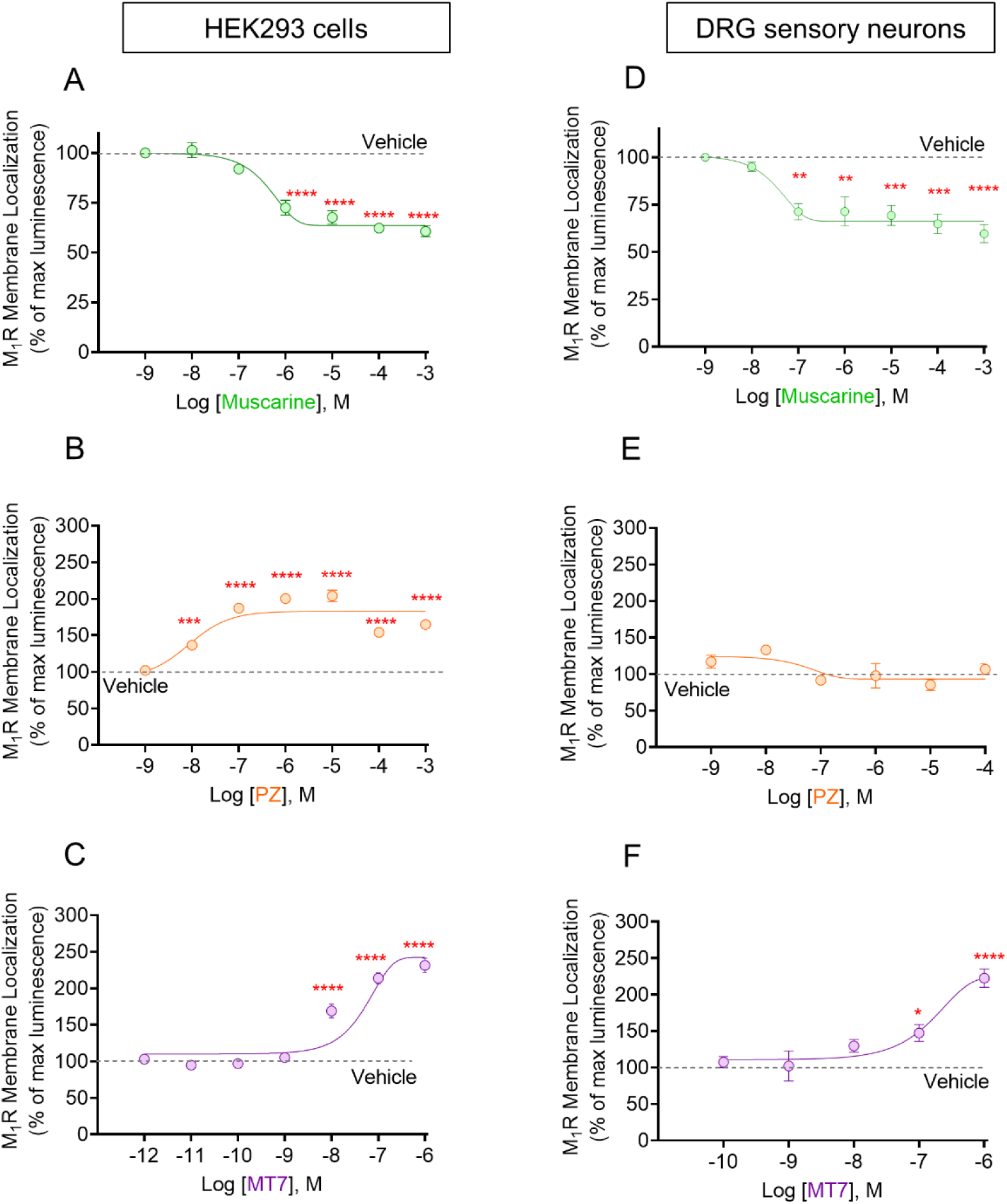
Pirenzepine and MT7 enhance accumulation of M_1_R at the surface of HEK293 cells DRG neurons. Dose response curve for receptor internalization performed for muscarine (**A&D**, 60 min), pirenzepine (**B&E**, 60 min), and MT7 (**C&F**, 60 min) in both HEK293 cells and DRG sensory neurons. Data show mean ± SEM (1-way ANOVA with Dunnett’s post-hoc test). Comparison to vehicle-treated group; ****P<0.0001, ***P<0.001, **P<0.01 and *P<0.05, n=4.

### β-arrestins (but not GRKs) are necessary for ERK activation by PZ and MT7

We investigated the factors that might be involved in the β-arrestin-mediated signaling and ERK activation by PZ and MT7 in M_1_R-transfected HEK293 cells. First, we examined the role of Gα_q_ protein activity on the PZ/MT7-induced ERK activation. Results showed that inhibition of Gα_q_ protein activity by YM-254890 had no effect on PZ/MT7-induced ERK activation in HEK293 (Fig. 8A&B). However, genetic ablation of β-arrestin1/2 abolished ERK activation induced by PZ/MT7 (Fig. 8C&D) and confirming our previous work (*24*). Surprisingly, ablation of GRKs (2, 3, 5, and 6) failed to suppress ERK activation in response to PZ/MT7 (Fig. 8E&F). It is interesting that treating non-transfected HEK293 cells with CMPD101, a GRK2/3 inhibitor, decreased ERK activation (Fig. S5), while the same treatment increased ERK activation in M_1_R-transfected HEK293 (Fig. 8G&H). Treating M_1_R-transfected HEK293 cells with CMPD101 significantly increased ERK activation in response to PZ/MT7 (Fig. 8G&H).

**Fig. 8.**
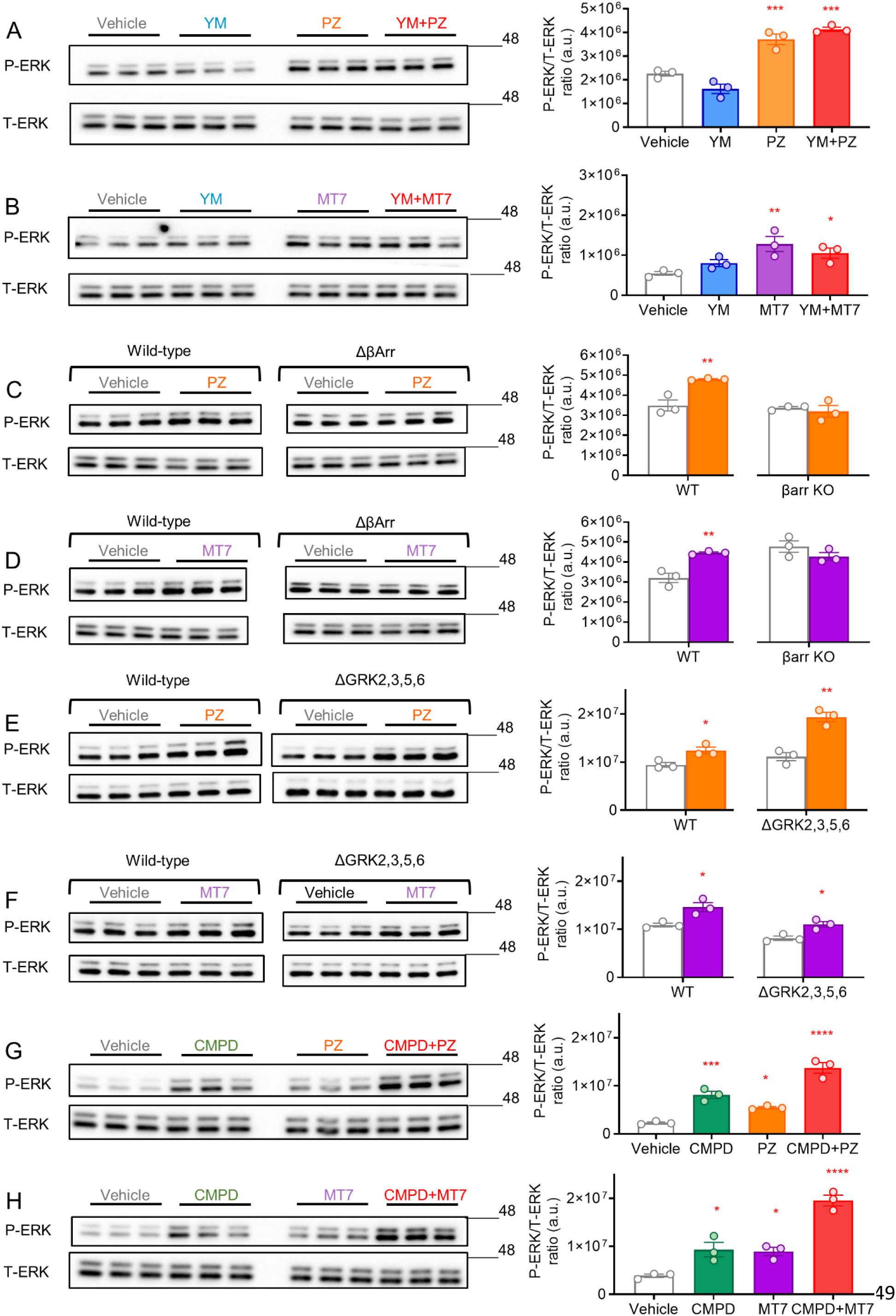
β-arrestins (but not GRKs and Gq-protein) are necessary for ERK phosphorylation following M_1_R antagonist treatment. (A&B) M_1_R-transfected HEK293 cells were pre-treated with vehicle or YM-254890 (1 µM, 60 min) before treatment with **(A)** pirenzepine (PZ, 1 µM, 60 min) or **(B)** MT7 (100 nM, 60 min). Immunoblotting for T-ERK and P-ERK proteins was conducted. Data represent P-ERK/T-ERK ratio (a.u.). Data are reported as the mean ± SEM (1-way ANOVA with Dunnett’s post-hoc test). Comparison to vehicle-treated group; ***P<0.001, **P<0.01, and *P<0.05, n=3. **(C-F)** hM_1_R transfected HEK293 cells (WT, ΔβArrestin1/2 or ΔGRK2,3,5,6) were treated with **(C&E)** pirenzepine (1 µM, 60 min) or **(D&F)** MT7 (100 nM, 60 min). Data are reported as the mean ± SEM. *t*-test was used for comparing vehicle-treated and drug-treated groups; **P<0.01 and *P<0.05, n=3. **(G&H)** hM_1_R transfected HEK293 cells were pretreated with vehicle/CMPD101 (30 µM, 10 min), and then received **(G)** PZ (1 µM, 60 min), and **(H)** MT7 (100 nM, 60 min). Data are reported as the mean ± SEM (1-way ANOVA with Dunnett’s post-hoc test). Comparison to vehicle-treated group; ****P<0.0001, ***P<0.001, and *P<0.05, n=3.

### CK2 inhibition prevented PZ-induced ERK activation and neurite outgrowth in DRG neurons

Fig. 8E&F demonstrated that GRKs (2, 3, 5, and 6) had no role in ERK activation mediated by PZ/MT7. Hence, alternative kinases such as CK1 and CK2 might be responsible for M_1_R phosphorylation, as previously reported (*4, 31, 35*). We examined the role of CK2 based on the previously reported work indicating that CK2 inhibition blocks M_1_R-mediated ERK activity in response to agonists (*4*). As seen in Fig. 9A, BRET results show that 4,5,6,7-tetrabromo-2-azabenzimidazole (TBB; a well characterized CK2 inhibitor (*36*), 1 µM, 10 min) pretreatment significantly reduced β-arrestin2 recruitment to M_1_R by PZ (1 µM, 30 min). Similarly, TBB pretreatment blocked PZ-induced ERK activation in HEK293 cells (Fig. 9B) and DRG neurons (Fig. 9C). Further, knockdown of *Csnk2a1* and CK2 depletion (Fig. S6) significantly reduced PZ-induced ERK activation in HEK293 cells (Fig. 9D). PZ (1 µM, 42 h) significantly elevated total neurite outgrowth in DRG neurons, and co-administration of TBB (30 µM) attenuated the positive effect of PZ (Fig. 9E). This result revealed for the first time that PZ-dependent blockade of M_1_R operated via CK2 to drive neurite outgrowth in primary adult neurons.

**Fig. 9.**
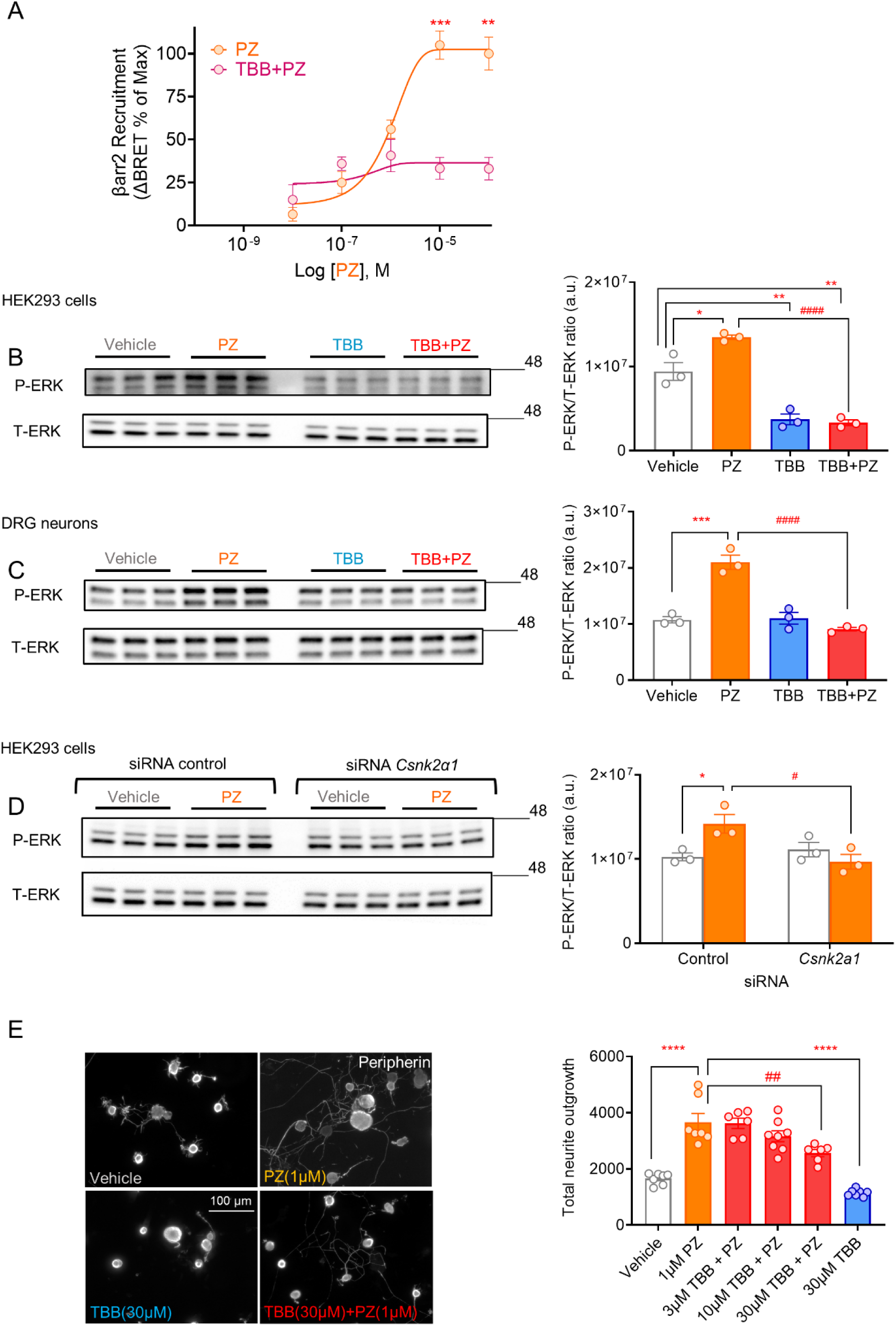
CK2 inhibition attenuated β-arrestin recruitment, ERK activation and neurite outgrowth by pirenzepine in DRG sensory neurons. **(A)** TBB (1 µM, 10 min) pretreatment blocked pirenzepine-induced β-arrestin2 recruitment to hM_1_R in HEK293 cells. Values are expressed as the mean ± SEM, and were analyzed using *t*-test. Comparison between PZ (100 µM) and TBB + PZ (100 µM); ***P<0.001 and comparison between PZ (10 µM) and TBB + PZ (10 µM); **P<0.01, n=4. **(B)** M_1_R-transfected HEK293 cells and **(C)** DRG neurons were pretreated with TBB (1 µM or 10 µM, 10 min, respectively), and then received pirenzepine (PZ, 1 µM, 60 min). Data are reported as the mean ± SEM (1-way ANOVA with Tukey’s post-hoc test). Comparison to vehicle; ***P<0.001, **P<0.01 and *P<0.05. Comparison to PZ-treated group; ####P<0.0001, n=3. **(D)** HEK293 cells were co-transfected with 25 nM of control or *Csnk2a1* siRNA and hM_1_R, and then received vehicle or pirenzepine (PZ, 1 µM, 60 min). Values are expressed as the mean ± SEM (2-way ANOVA with Tukey’s post-hoc test). Comparison between vehicle and PZ-treated cells in siRNA control group; *P<0.05, and comparison between PZ-treated cells in siRNA control and *Csnk2a1* groups; #P<0.05, n=3. **(E)** DRG sensory neurons were treated with pirenzepine (PZ,1 µM), and different doses of TBB in combination with (PZ, 1 µM) for 42 h. Cells were fixed and immune-stained for peripherin, and total neurite outgrowth was assessed. Values are means ± SEM (1-way ANOVA with Tukey’s post hoc test). Comparison between vehicle/TBB 30 µM-treated cells with PZ (1 µM)-treated group; ****P<0.0001, and comparison between PZ (1 µM)-treated cells and PZ (1 µM) + TBB (30 µM) combination group; ##P<0.01, n = 6-8 replicate cultures.

## DISCUSSION

We provide evidence that PZ and MT7 act as β-arrestin-biased agonists at M_1_R by recruiting β-arrestin2 and associated signaling pathways in preference over G protein activation in HEK293 cells and adult primary DRG sensory neurons. The data demonstrates that phosphorylation of M_1_R at specific ICL3 residues is necessary for β-arrestin-mediated ERK activation. ERK activation by PZ/MT7 required β-arrestins and activation of CK2 (but not GRKs and Gα_q_ protein). Lastly, CK2 inhibition blocked PZ-dependent enhancement of neurite outgrowth in DRG sensory neurons. The pharmacological profile of PZ and MT7 is consistent with previously reported β-arrestin biased agonists, such as carvedilol, including lack of G protein agonism, altered receptor phosphorylation and β-arrestin-dependent signaling (*26, 37*).

Acute treatment (up to 5 min) with PZ and MT7, two structurally and functionally distinct M_1_R ligands, exhibited antagonistic effects in response to muscarine/carbachol and blocked M_1_R-arrestin interaction. However, longer treatment (e.g. 30 min) of PZ or MT7 in the absence of full agonist effectively recruited β-arrestin2 (Fig. 2D & E, Fig. 3E & F) and induced ERK activation (Fig. 2F&G, Fig. 3E&F). These pharmacological effects were markedly different from the orthosteric agonist carbachol/muscarine with much less ability to promptly recruit β-arrestin2 and initiate receptor internalization. PZ/MT7 time-dependently activated the β-arrestin-ERK pathway while inducing no G protein signaling and receptor internalization. This data suggests these ligands are capable of inducing specific receptor conformations and signaling as partial and biased agonists: acting as antagonists in the presence of a full unbiased agonist. In this regard, studies using transfected model systems showed PZ upregulated M_1_R, induced stabilization of dimers/oligomers at cell surface (*38, 39*), and resulted in slower diffusion of receptor, which might reduce the possibility of interactions between receptor and its cognate G proteins (*40*). Similarly, MT7 was reported to bind to the dimerized form of M_1_R and stabilized this receptor state at the cell surface (*41, 42*). Indeed, our results reveal that PZ/MT7 upregulated M_1_R at the surface of HEK293 cells and DRG sensory neurons for up to 3 h (Fig. 7 and Fig. S4). Other studies reported that ERK activation by β-arrestin-biased signaling at muscarinic receptors did not necessarily internalize the receptor (*4, 43*). In line with previous reports, we showed M_1_R expression in different sub-types of DRG sensory neurons (*23, 44*), and only M_1_R-positive neurons exhibited ERK activation in response to MT7. We recently demonstrated that M_1_R overexpression markedly decreased neurite outgrowth and mitochondrial function in sensory neurons, and ERK activity played a major role in the neuritogenic effects of MT7 (*16, 24*).

Binding of β-arrestins to phosphorylated GPCRs regulates many aspects of cellular signaling. A significant body of evidence supports the “barcode hypothesis”, which explains how different phosphorylation of residues at GPCRs results in diverse β-arrestin-associated signaling responses (*25, 31, 45*). A previous study by Butcher *et al*. revealed that acetylcholine treatment phosphorylated M_1_R at 12 residues at ICL3 and 2 residues at the C-terminus (*13*). In comparison to acetylcholine, our results showed that PZ phosphorylated hM_1_R at 6 positions (similar to (*13*)) at ICL3 (Fig. 5A), and subsequent mutation at these residues (deletion of all 6 positions or mutation at Ser^251^ and Thr^254^) significantly reduced β-arrestin2 recruitment to M_1_R and abolished ERK activation. In the case of MT7, deletion of all phosphorylated residues diminished β-arrestin2 recruitment and ERK activation, but mutation at Ser^251^ and Thr^254^ had no effect. This suggests that phosphorylation of a specific set of ICL3 residues is required for PZ- or MT7-induced β-arrestin biased activation of ERK. Surprisingly, ICL3 mutations had no effect on muscarine-induced β-arrestin2 recruitment, and even increased ERK activation. These results indicate that each ligand has a specific signature with respect to the phosphorylation pattern generated in the ICL3 of M_1_R.

What factors might contribute to the β-arrestin biased activation of ERK by PZ/MT7? First, Gα_q_ inhibition did not alter the PZ/MT7-induced ERK activity (Fig. 8A & B). Recent studies demonstrated that ERK activity requires active G proteins, including ERK activation by arrestins (*46, 47*). Our data reveals that PZ/MT7 did not activate Gα_q_ signaling, and Gα_q_ inhibition did not modify the effects of PZ/MT7 on ERK activation. However, the effect of inhibition of alternative G proteins on putative ERK activity induced by PZ/MT7 was not tested. Furthermore, the data demonstrates that arrestins are necessary for PZ/MT7-induced ERK activity as lack of arrestins abolished the effect of these drugs. Other studies on β-arrestin biased ligands reported the same observation where arrestins play a major role in the biased effects of these ligands (*26, 37, 48*). Surprisingly, pharmacological inhibition of GRK2/3 with CMPD101 led to an increase in ERK activity mediated by PZ/MT7 (Fig. 8G & H), which indicates other kinases might be involved in M_1_R phosphorylation. Recent work by Drube and colleagues showed that no specific GRK is assigned for M_1_R recruitment of arrestins (*35*). Indeed, casein kinases phosphorylate muscarinic receptors (particularly M_1_R and M_3_R) (*4, 5, 49*), and among them, CK2 contributes to ERK activation by M_1_R (*4*). Our findings showed siRNA-based or pharmacological CK2 inhibition strikingly reduced ERK activation by PZ in HEK293 cells.

In DRG sensory neurons, the PZ-induced activation of ERK was dependent on CK2 (Fig. 9). Previous work by Jung *et al*., showed that CK2 inhibition abolished arrestin-mediated ERK activity by M_1_R agonists (*4*). CK2 inhibition had no effect on basal ERK activity in DRG neurons, but significantly suppressed ERK activity in HEK293 cells, which explains drug-response differences between cell types. Indeed, CK2 is expressed in DRG sensory neurons, and plays a major role in neurite outgrowth where knockdown of CK2 suppressed neurite outgrowth (*50, 51*). Intriguingly, our results showed that CK2 inhibition blocked PZ-induced neurite outgrowth in cultured DRG sensory neurons. These findings suggest that CK2 is necessary for arrestin-mediated ERK activity by PZ and suppressing ERK activity through CK2 inhibition blocks PZ-induced neurite outgrowth in sensory neurons.

Differences in signal transduction processes between PZ (as an orthosteric antagonist) and MT7 (as a NAM) were noted in this work. Unlike PZ, MT7 is a cell impermeable large molecule with a very slow dissociation rate, which determines its primary action is at the cell surface (*30, 52*). Our findings showed that MT7 is more potent and rapid than PZ in inducing ERK activity and M_1_R upregulation at the cell surface. Also, MT7 showed no sensitivity to Ser^251^-Thr^254^ mutation in M_1_R and its induction of β-arrestin2-ERK signaling was unaffected (Fig. 6). These discrepancies may be related to several factors such as differences in molecular size, binding location at M_1_R, dissociation rate, and permeability of these two ligands.

Our study has some limitations. We used two model systems, HEK293 cells and adult DRG sensory neurons, and overexpressed modified proteins in these cells, which might not mimic real physiological conditions of sensory neurons. In addition, DRG sensory neurons are classified into several sub-types based on their soma size, expression of specific markers, and their conduction properties (*53*). Consequently, we showed M_1_R expression (using specific MT7-ATTO590 marker) on DRG neurons with varying phenotypes, but did not delineate which sub-type of DRG neuron was responding to PZ/MT7 with β-arrestin biased activation of ERK.

Our future studies could focus on the cross-talk between M_1_R and CK2 in different *in vivo* models of peripheral neuropathy, and uncover the molecular pathways associated with M_1_R biased activity in different types of DRG neurons to find an appropriate treatment for peripheral neuropathy with minimal side effects. In summary, we showed that antimuscarinic drugs, PZ and MT7, activated M_1_R towards the β-arrestin signaling pathway in both HEK293 and DRG sensory neurons. These drugs acted as β-arrestin-biased agonists and stimulated neurite outgrowth and could be potential treatment options for preventing/reversing nerve degeneration in different types of peripheral neuropathy.

## MATERIALS AND METHODS

### Chemicals

Muscarine, carbachol, atropine, and PZ were purchased from MilliporeSigma Canada (Oakville, ON, Canada). MT7, fluorescent dye ATTO Fluor 590-conjugated MT7 (MT7-ATTO590), and YM-254890 were purchased from Alomone Labs (Jerusalem, Israel) and Cayman Chemicals (Ann Arbor, MI, USA), respectively. CMPD101 and TBB were purchased from Tocris (Minneapolis, MN, USA).

### Cell Culture and transfection

#### Cell lines

Wild-type (WT) Human Embryonic Kidney (HEK293) cells were grown in Dulbecco’s modified Eagle’s medium (DMEM)/F12 (Thermo Fisher Scientific, Waltham, MA, USA) supplemented with 5% heat inactivated fetal bovine serum (FBS, Thermo Fisher Scientific, Waltham, MA, USA) and 1% penicillin and streptomycin (MilliporeSigma Canada, Oakville, ON, Canada) at 37°C and 5% CO_2_. The CRISPR/Cas9 mediated β-arrestin 1/2 knockout (KO) (Δβarr1/2) and GRK2, 3, 5, and 6 KO (ΔGRK2,3,5,6) HEK293 cell lines were designed and generated by Dr. Inoue laboratory (*54, 55*). Lipofectamine 3000 reagent (Thermo Fisher Scientific, Waltham, MA, USA) was used to transfect the cells (details are provided in each experimental section). For siRNA transfection in HEK293 cells, 25 nM siRNA control (ON-TARGETplus Non-targeting siRNA, 5’UGGUUUACAUGUCGACUAA 3’, Dharmacon, Cat#D-001810-01-05) and siRNA against CK2 (On-TARGETplus-SMART pool siRNA against *Csnk2a1*, Dharmacon, Cat#L-096197-02-0005) were used with Lipofectamine 3000 reagent and were incubated for 48 h.

#### Adult rat DRG sensory neuron culture

Adult male Sprague-Dawley rats, adult male wild-type (C57BL/6) and M_1_R-KO (C57BL/6 background, line1784; Taconic Biosciences Inc., Hudson, NY, USA) were euthanized and DRG were dissected, dissociated, and cultured as previously described (*56*). All animal procedures followed the guidelines of the University of Manitoba Animal Care Committee using the Canadian Committee on Animal Care (CCAC) rules. In brief, DRG neurons were cultured in Ham’s F-12 media (Wisent Inc., Saint-Jean Baptiste, QC, Canada) supplemented with modified Bottenstein’s N2 additives (0.1 mg/ml transferrin, 20 nM progesterone, 100 mM putrescine, 30 nM sodium selenite, 0.1 mg/ml BSA, all purchased from MilliporeSigma Canada, Oakville, ON, Canada). Media was also supplemented with 0.146 g/L L-glutamine (Thermo Fisher Scientific, Waltham, MA), a low dose cocktail of neurotrophic factors (0.1 ng/ml NGF, 1.0 ng/ml GDNF and 1 ng/ml NT-3, all from Thermo Fisher Scientific, Waltham, MA USA), 0.1 nM insulin and 1X antibiotic antimycotic solution (MilliporeSigma Canada, Oakville, ON, Canada). DRG neurons were transfected using NeuroMag (Oz Biosciences, San Diego, CA, USA) by magnetofection method (details are provided in each experimental section).

### Plasmid construction

NLuc-M_1_R, HaloTag-β-arrestin2, Halo-M_1_R, and HiBiT-M_1_R plasmids were purchased from GeneCopoeia Inc. (Rockville, MD, USA). All gene synthesis, cloning and mutagenesis for NLuc-M_1_R was by GenScript (Piscataway, NJ, USA) and confirmed by DNA sequencing. Phospho-proteomics results were used to design M_1_R mutant plasmids. Mutant M_1_R plasmids used in this study were 1) NLuc-M_1_R with Thr^230^, Ser^251^, Thr^254^, Ser^321^, Thr^354^, and Ser^356^ deletion at ICL3 loop (M_1_RΔICL3), and 2) NLuc-M_1_R with Ser^251^ and Thr^254^ substitution with Alanine at ICL3 loop (M_1_R-S251A, T254A). pcDNA3.1 and -22F cAMP pGloSensor.

### β-arrestin recruitment

#### HEK293 cells

For BRET-based examination of β-arrestin2 recruitment to M_1_R in HEK293 cells, 400000 cells were seeded onto 6-well plates per well. On the following day, cells were co-transfected using Lipofectamine 3000 with NLuc-hM_1_R/mutant NLuc-hM_1_R plasmids/pcDNA3.1 (20 ng) and HaloTag-β-arrestin2 (800 ng) per well (1:40 ratio), following the manufacturer’s protocol. Calculation of the ratio and plasmid concentrations were obtained from saturation assay studies in the lab. After 24 h, cells were detached, centrifuged, and resuspended in OPTI-MEM media (no phenol red, with 4% FBS) plus HaloTag® NanoBRET™ 618 Ligand (Promega, Madison, WI, USA, final concentration 100 nM), and seeded (100 µl/well containing 25000 cells) into poly-D-L-ornithine hydrobromide (PDO)-coated 96-well white plates (OPTI-MEM from Thermo Fisher Scientific, Waltham, MA and PDO is from MilliporeSigma Canada, Oakville, ON, Canada). After 20 h, cell media was replaced with 90 µl HBSS (Thermo Fisher, Waltham, MA, USA) and 10 µl of 10X drugs. In the case of DRG neurons, 7500 cells were seeded into PDO and laminin-coated 96-well white plates and left unharmed for 36 h. DRG neurons were co-transfected with NLuc-hM_1_R/pcDNA3.1 (50 ng) and HaloTag-β-arrestin2 (200 ng) per well (1:4 ratio) by magnetofection using NeuroMag (Oz Biosciences, San Diego, CA, USA) according to the manufacturer’s protocol. Forty-eight hours after transfection, HaloTag® NanoBRET™ 618 Ligand (Promega, Madison, WI, USA, final concentration 100 nM) was added to wells, and then after 20 hours cell media was replaced with 90 µl of Ham’s F-12 media (no phenol red) for 15 min, and then 10 µl of 10X drugs were added to wells. After incubation with drugs (5 min for carbachol/muscarine, and 30 min for PZ/MT7), 25 µl of Nano-Glo substrate (Promega, Madison, WI, USA, 1:400 final dilution) was added to wells, and a Synergy Neo2 plate reader (Biotek/Agilent Technologies, Santa Clara, CA, USA) was used to read luminescence and fluorescence to obtain the BRET ratio (610nm/460nm). β-arrestin2 recruitment was calculated as the change in BRET ratio (ΔBRET) following treatments and normalization to the baseline and test compound/vehicle response.

### IP1 assay

As a measure of GPCR-associated Gα_q_ signaling intracellular IP1 levels were measured using the IP-One assay (Cisbio Bioassays, Waltham, MA, USA). DRG sensory neurons were plated onto PDO and laminin-coated 384-well white microplates (5000 cells per well). HEK293 cells transiently transfected with plasmids containing the hM_1_R-WT/pcDNA3.1, and were plated onto 384-well white microplates (20000 cells per well) and were treated with stimulation buffer with or without 1) different doses of M_1_R agonist (muscarine), 2) different doses of M_1_R antagonists (PZ and MT7) or 3) both M_1_R antagonist (different doses of PZ/MT7) and agonist (muscarine, 1 mM) for different time points at 37 °C. Cells were lysed by addition of the supplied buffer containing d2-labeled IP1, followed by addition of terbium cryptate-labeled anti-IP1 antibody, according to the manufacturer’s instructions. Plates were incubated for 1 h at room temperature and fluorescence signals were measured at 620 and 665 nm using the Biotek Synergy Neo2 plate reader. HTRF ratio 665 nm/620 nm of each well were extrapolated on the standard curve to obtain IP1 concentrations.

### Glosensor cAMP assay

HEK293 cells (400000 cells/well, 6-well plate) transiently transfected with plasmids containing the hM_1_R-WT (180 ng) and -22F cAMP pGloSensor (1080 ng) using Lipofectamine 3000. After 24 h, cells were detached, washed, centrifuged, resuspended in HBSS, and seeded (65µl/well containing 40000 cells) into 96-well white plates. Then, cells were incubated with Glosensor reagent (Promega, Madison, WI, USA, 10 µl/well) for two hours in the dark. Cells were pre-treated with 1) various concentrations (25 µl of 4X drug) of muscarine, PZ, and MT7, and 2) muscarine (1 mM) + various doses of PZ/MT7 and baseline luminescence was read on a Synergy Neo2 plate reader. After 5 min, cells were treated with 300 nM of forskolin (FSK, MilliporeSigma Canada, Oakville, ON, Canada)) and readings were continued for up to 20 min. Effects of drugs on M_1_R-induced cAMP changes was calculated as the decrease/increase in FSK-induced luminescence amplitude.

### Western blotting

HEK293 cells or DRG sensory neurons were lysed and homogenized in ice-cold RIPA buffer containing protease and phosphatase inhibitor cocktail (ThermoFisher Scientific, Waltham, MA, USA). DC protein assay (Bio-Rad, Hercules, CA, USA) was used for protein assay, and Western blot analysis was performed. Protein samples (20 µg total protein/lane for HEK293 cells and 5 µg total protein/lane for DRG neurons) were resolved and separated by 10% SDS-PAGE, and transferred to a nitrocellulose membrane (Bio-Rad, Hercules, CA, USA) using Trans-Blot Turbo Transfer System (Bio-Rad, Hercules, CA, USA) and immunoblotted with specific antibodies against total ERK (T-ERK; 1:3000, Santa Cruz Biotechnology,Dallas, TX, USA, sc-514302), and phosphor-ERK (P-ERK, 1:3000, Cell Signaling Technology, Danvers, MA, USA, Cat# #9101). HRP-conjugated goat anti-rabbit IgG or goat anti-mouse IgG (Jackson ImmunoResearch Laboratories, West Grove, PA, USA) were used for secondary antibodies. The blots were incubated in Clarity™ Western ECL substrate (Bio-Rad, Hercules, CA, USA) or SignalFire™ ECL Reagent (Cell Signaling Technology, Danvers, MA, USA) and imaged using a Bio-Rad ChemiDoc image analyzer (Bio-Rad, Hercules, CA, USA).

### Mass spectrometry

#### Purification of hM_1_R

For hM_1_R purification, HaloTag® Mammalian Protein Purification System (Promega, Madison, WI, USA) was used. In brief, HEK293 cells (five 15-cm plates) were transfected with Halo-tagged hM_1_R construct (20 µg/dish) using Lipofectamine 3000 for 24 h. Cells were washed twice with PBS, were incubated in serum-free DMEM for 6 h, and then media were removed and replaced with fresh serum-free DMEM containing 1 µM PZ for 30 min. Cells were scraped into 5 ml of lysis buffer (50 mM Tris-HCl, pH 7.5, 150 mM NaCl, 10 mM NaF, (all from MillipoeSigma Canada, Oakville, ON, Canada) and protease inhibitor cocktail (Promega, Madison, WI, USA) and lysed by sonication on ice. Cellular debris was cleared by centrifugation, and 75 µl of HaloLink™ Resin (Promega, Madison, WI, USA), equilibrated with TBS containing 10XCMC or 0.087% n-Dodecyl-β-D Maltopyranoside (DDM) (Thermo Fisher Scientific, Waltham, MA, USA) was added. The samples were incubated overnight at 4 °C with rotation, and then supernatant was discarded and resin was washed (three times) with TBS+DDM10XCMC and TBS+DDM 3XCMC, respectively. The resin was treated with elution buffer (150 mM NaCl, 50 mM Tris-HCl (pH=7.5) + DTT (MilliporeSigma Canada, Oakville, ON, Canada, 1mM) + NaF (10mM) + DDM 3XCMC) containing HaloTEV Protease (Promega, Madison, WI, USA) overnight at 4 °C with rotation. Resin was cleared by centrifugation, and purified samples were collected and sent to the Manitoba Centre for Proteomics and Systems Biology (MCPSB) and to the University of Virginia Biomedical Research Facility for LC/MS/MS analysis. The following LC-MS/MS analysis methodology is for the Manitoba Centre for Proteomics and Systems Biology (MCPSB). The methodology for the University of Virginia Biomedical Research Facility is in the supplementary materials.

#### LC-MS/MS analysis

Per sample, 10 μg of protein was separated by SDS-PAGE. The gel was fixed with 5% acetic acid in 50% methanol for 30 minutes and stained with GelCode Blue Safe Protein Stain (ThermoFisher Scientific, catalog # 24596) overnight, followed by destaining in water. The 50 kDa protein bands were excised and gel pieces were destained three times with 500 µL of acetonitrile and vortexed for 10 minutes (repeated twice). Gel pieces were dried in a speedvac, followed by incubation with 10 mM dithiothreitol (DTT) in 25 mM ammonium bicarbonate (ABC) at 56 °C for 30 minutes to reduce disulfide bonds. Gel pieces were dehydrated in 500 µL acetonitrile for 10 minutes, followed by incubation with 55 mM iodoacetamide in 25 mM ABC for 30 min at RT in the dark, to alkylate free Cys residues. Gel pieces were dehydrated with 500 µL acetonitrile for 10 minutes, and access iodoacetamide was quenched with 10 mM DTT in 25 mM ABC for 10 minutes at RT. Gel pieces were dehydrated in 500 µL acetonitrile for 10 minutes and rehydrated with digestion buffer containing 12.5 ng/µL trypsin/LysC (Promega) in 50 mM ABC. Proteins were digested at 37 °C overnight. Supernatants were collected and combined with four extracts collected after successively incubating the gel pieces once with H_2_O and three times with 5% formic acid in 50% acetonitrile. Sample volumes were reduced in a speedvac, and generated peptides were desalted using C18 solid phase extraction tips (ThermoFisher Scientific, catalog #87782). Per sample, 60% of the peptide extract was analyzed by nano-LC-MS/MS on an Orbitrap Exploris 480 (Thermo Fisher Scientific, Bremen, Germany) online-coupled to an Easy-nLC 1200 (Thermo Fisher Scientific). Mobile phase A was 0.1% (v/v) formic acid and mobile phase B was 0.1% (v/v) formic acid in 80% acetonitrile (LC-MS grade). Peptides were separated on a Luna C18(2) column (3 μm particle size; Phenomenex, Torrance, CA) packed in-house in a Pico-Frit (100 μm x 30 cm) capillary (New Objective, Woburn, MA) with the following gradient at a flow rate of 300 nL/min: 2-6% B in 3 minutes, 6-30% B in 36 min, 30-50% in 5 min, 50-90% in 1 min, remain at 90% for 5 min. The Orbitrap Exploris 480 was ran in data-dependent acquisition (DDA) mode. Survey scans were acquired from m/z 380-1500 at a resolution of 45,000 (at m/z 200), with a maximum ion injection time of 45 milliseconds, and a normalized automatic gain control (AGC) target of 200%. The top 12 precursor ions with charge states 2-5+ were isolated with a m/z 1.6 window, fragmented with a normalized collision energy of 30%, and MS/MS scans were acquired at a resolution of 30,000. AGC target value was set to 250%, with a maximum ion injection time set to auto.

Mass spectrometric data were searched using Proteome Discoverer v3.0 using SequestHT, MS Amanda 2.0, Percolator, and ptmRS nodes, against a human Swissprot protein database (downloaded from Uniprot.org in July 2022, 20399 target sequences) with the following search settings: trypsin as enzyme with a maximum of 2 missed cleavages, oxidation of Met (+15.995 Da), phosphorylation of Ser/Thr/Tyr (+79.966 Da) and deamidation of Asn/Gln (+0.984 Da) as variable and carbamidomethylation of Cys (+57.021 Da) as static modifications. MS and MS/MS tolerances were set to 10 ppm and 0.02 Da, respectively. The data was filtered to 1% false discovery rate on the protein, peptide, and peptide-spectrum-match levels, and only phosphopeptides with ptmRS site localization probabilities ≥99% were considered.

### Internalization assay

Human M_1_R internalization was measured by HiBiT-based receptor internalization assay (Promega). HEK293 cells were transiently transfected with hM_1_R-HiBiT plasmid (100 ng) and pcDNA3.1 (9900 ng) in suspension (OPTI-MEM no phenol red) using Lipofectamine ® 3000. Then, cells were seeded (95 µl/25000 cells) into PDO-coated white 96-well plates. After 20 h, cells were treated with 10 µl of (10X) different doses of muscarine, PZ, and MT7 for 1 h, and were incubated at 37^0^C + 5% CO_2_ for 1 h. Then 100 µl of 2× substrate buffer consisting of 1:100 of a LgBiT stock solution and 1:50 dilution of Nano-Glo® HiBiT Extracellular Substrate in the OPTI-MEM (no phenol red) was added to wells, and a Synergy Neo2 plate reader was used to read luminescence. Luminescence signals were averaged and normalized to the vehicle stimulation count.

For the DRG sensory neurons, 7500 cells were seeded into PDO and laminin-coated 96-well white plates and left unharmed for 36 h. DRG neurons were transfected with hM_1_R-HiBiT plasmid (25 ng) per well by magnetofection using NeuroMag according to the manufacturer’s protocol. Forty-eight hours after transfection, media was removed and replaced with 90 µl of Ham’s F-12 media (no phenol red) supplemented with N2 additives, and cells were incubated at 37^0^C + 5% CO_2_ for 15 min. Then, 10 µl of (10X) different doses of muscarine, PZ, and MT7 for different time-points. Finally, 100 µl of 2× substrate buffer consisting of 1:100 of a LgBiT stock solution (Promega) and 1:50 dilution of Nano-Glo® HiBiT Extracellular Substrate in the Ham’s F-12 media (no phenol red) was added to wells, and a Synergy Neo2 plate reader was used to read luminescence. Luminescence signals were averaged and normalized to the vehicle stimulation count.

### Immunohistochemistry (IHC) and immunocytochemistry (ICC)

For the IHC, mouse DRG tissues derived from C57BL/6 mice (*56*) were exposed to MT7-ATTO590 (100 nM, overnight) at 37 °C CO_2_ incubator, and then were fixed in PFA 2% (Thermo Fisher Scientic, Waltham, MA, USA), cryoprotected in 20% sucrose, and finally embedded in Tissue-Tek O.C.T. (Sakura Finetek USA, Torrence, CA, USA) to prepare 7 μm tissue sections. All sections were incubated with β-tubulin III antibody (1:500; T8578, MilliporeSigma Canada, Oakville, ON, Canada) overnight at 4 °C, and then stained for 1 h with Alexa Fluor 488-conjugated anti-mouse IgG (1:1000; Thermo Fisher Scientific, Waltham, MA, USA) at room temperature. For the ICC mouse DRG neurons were cultured on glass coverslips, incubated with MT7-ATTO590 (100 nM, 90 minutes) at 37° C in a CO_2_ incubator, and then were fixed with PFA 2%, and permeabilized with 0.3% Triton X-100 (MilliporeSigma Canada, Oakville, ON, Canada). Neurons were blocked by 5% BSA for 1 h, and then incubated with phospho-ERK antibody (P-ERK, 1:500, Cell Signaling Technology, Danvers, MA, USA Cat# #9101) overnight. Cells were washed with PBS (3X), and then stained for 1 h with Alexa Fluor 488-conjugated anti-rabbit IgG (1:1000; Thermo Fisher Scientific, Waltham, MA, USA) at room temperature. Coverslips were mounted on slides using VECTASHIELD antifade mounting medium with DAPI (Vectorlabs, Inc., Newark, CA, USA). All images were taken by using a Carl Zeiss LSM510 confocal or Axioscope-2 fluorescence microscope.

### Neurite outgrowth in adult DRG sensory neurons

DRG sensory neurons were cultured on glass coverslips and were treated with 1) vehicle 2) PZ (1 µM), 3) TBB (30 µM), or 4) combination of PZ (1 µM) + TBB (3, 10, and 30 µM). Appropriate doses for TBB were obtained from pilot studies in the lab. After 24 h, cells were fixed with 2% paraformaldehyde (pH 7.4, 15 min), and permeabilized with 0.3% Triton X-100 in PBS for 5 min. Cells were blocked by 10% blocking reagent (Roche Diagnostics, Laval, QC, Canada) and 20% FBS in PBS for 1 h, and then incubated with Peripherin antibody (1:1000, Santa Cruz Biotechnology, Dallas, TX, USA, sc-377093) overnight. Cells were washed with PBS (3X), and were incubated with Cy3-conjugated secondary antibody (Jackson ImmunoResearch, West Grove, PA, USA) for 1 h. Coverslips were mounted on slides using PERMAFLUOR mounting medium (Thermo Fisher Scientific, Waltham, MA, USA) and imaged using a Carl Zeiss Axioscope-2 fluorescence microscope equipped with equipped with an AxioCam camera. Neurite outgrowth was determined by collecting fluorescent signals as total pixel area for neurites and were analyzed by the high throughput NeurphologyJ plugin in ImageJ software after image enhancement. Total pixel area was normalized to number of cell bodies to calculate total neurite outgrowth per neuron.

### Statistical analysis

Excel (Microsoft, Redmond, WA, USA) and graphed in Prism 9.0 (GraphPad, San Diego, CA, USA) were used to analyze the data. Statistical tests were performed using a *t*-test, one or two-way ANOVA followed by multiple comparison’s tests. Details of statistical analysis and replicates are included in the figure legends. Lines represent the mean, and error bars signify the SEM, unless otherwise noted.

## ACKNOWLEDGMENTS

We thank Dr. Nicholas Sherman at the University of Virginia Biomedical Research Facility for supervising the initial LC-MS/MS analysis and bioinformatic work on phosphorylation sites of M_1_R in response to pirenzepine. We also thank Dr. Jürgen Wess, NIDDK, for the kind gift of the M_1_R KO mouse line. Graphical figures were created with BioRender.

## AUTHOR CONTRIBUTIONS

SA performed and designed all experiments (except where specified in the following paragraph). In addition, SA carried out data analysis and construction of figures. MA constructed and tested the M_1_R and β-arrestin2 plasmids for the BRET analysis. DRS performed the neurite outgrowth studies with cultured DRG sensory neurons. TMZW set up the use of fluorescently tagged MT7 and performed the related labelling studies in vitro and in sections of DRG. YL processed samples and performed mass spectroscopy for proteomic analysis. RPZ supervised the proteomic work and performed bioinformatic analysis. AI designed and generated the β-arrestin and GRK knockout HEK293 cells lines using CRISPR-Cas9 technology. HAD contributed to supervision of some methodology, data analysis, and design/construction of figures. SA wrote the 1^st^ draft of the manuscript which was then edited and refined by PF. All authors reviewed and edited the manuscript prior to submission. PF helped to design experiments, supervised all the work and obtained supporting funding.

## DATA AND MATERIALS AVAILABILITY

All data needed to evaluate the conclusions in the paper are present in the paper or the Supplementary Materials. All plasmids generated in this study will be distributed upon reasonable request and completion of a materials transfer agreement with University of Manitoba. All data reported in this paper will be shared by the lead contact upon request.

