## Supplementary data for "Antimuscarinic drugs exert β-arrestin-biased agonism at the muscarinic acetylcholine type 1 receptor"

**Supplementary materials**

**Supplementary figures: Figs. S1-6**

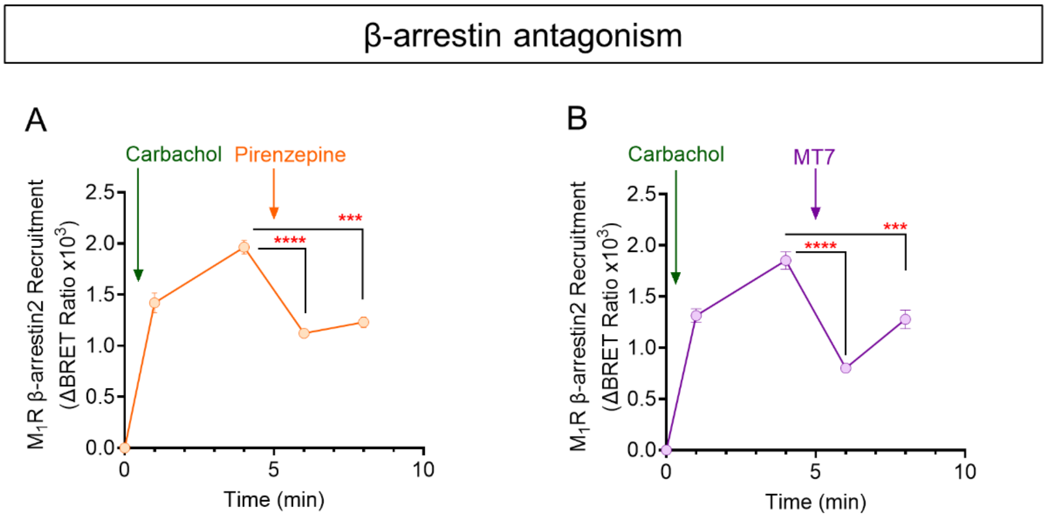


**Fig. S1. Antagonism of the carbachol-induced recruitment of HaloTag-β-arrestin2 to $hM_1R$.** Ligands were added at the times indicated. Carbachol (1 mM)-induced βArr2-Halo recruitment to $hM_1R$-Nluc was blocked by **(A)** pirenzepine (1 µM) and **(B)** MT7 (100 nM) in transiently co-transfected HEK293 cells. Data show mean ± SEM (1-way ANOVA with Dunnett's post-hoc test). ****P<0.0001, and ***P<0.001, n=3. Antagonist-treated groups were compared to carbachol-treated groups at minute 5.

A

SQPGAEGpS(251)PETPPGR

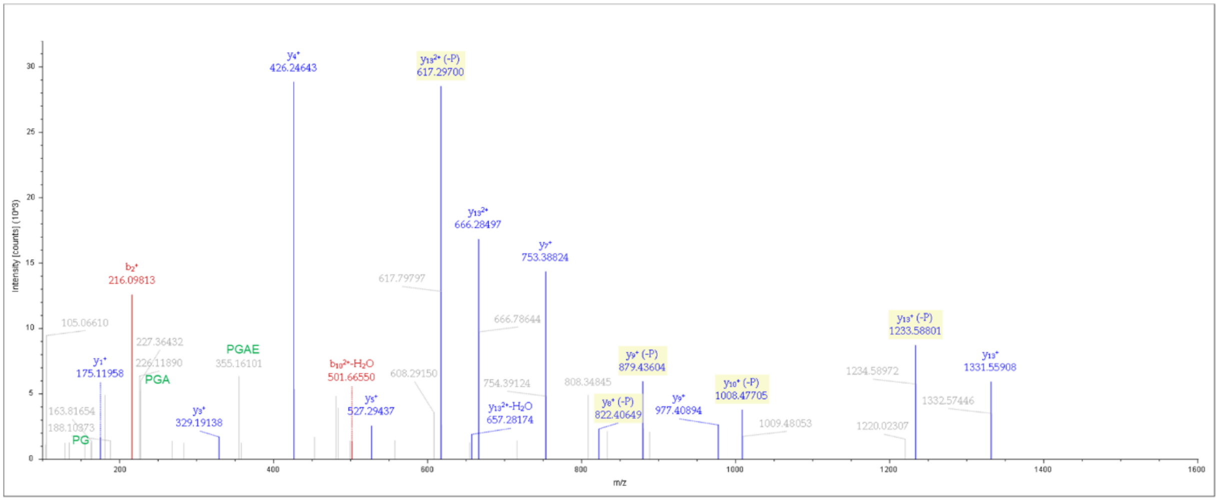


B

SQPGAEGSPEpT(254)PPGR

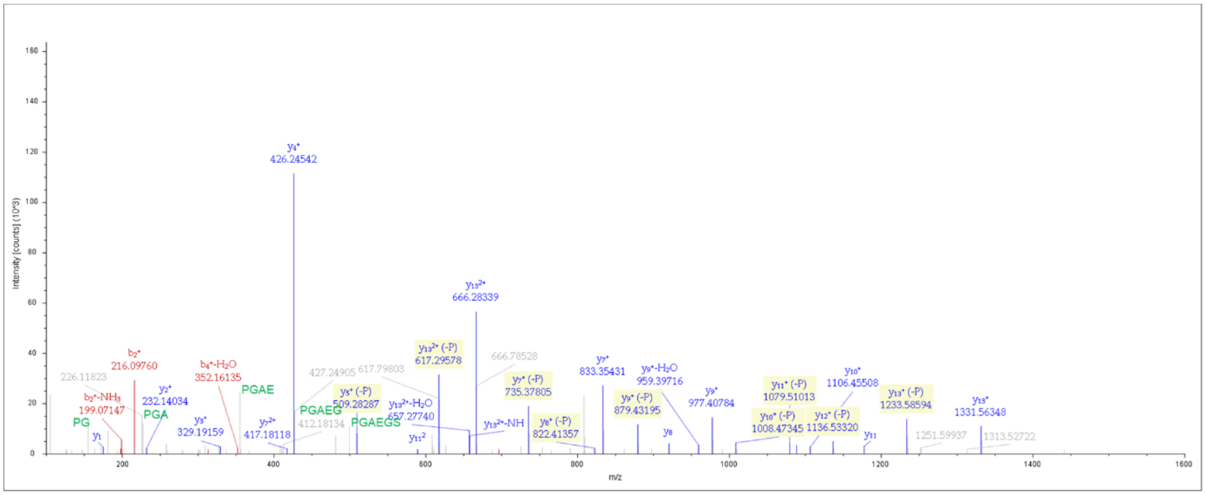


**Fig. S2. LC-MS/MS analysis identifies sites of hM$_1$R phosphorylation in HEK293 cells following pirenzepine treatment.** LC-MS/MS analysis performed by the Biomolecular Analysis Facility at University of Virginia Health System. **(A**) MS2 spectra of identified hM$_1$R phosphorylation at S251 in ICL3. **(B)** MS2 spectra of identified hM$_1$R phosphorylation at T254 in ICL3.

**Methodology for Fig. S2.** The gel pieces from the band were transferred to a siliconized tube and washed in 200 µL 50% methanol. The gel pieces were dehydrated in acetonitrile, rehydrated in 30 µL of 10 mM dithiolthreitol in 0.1 M ammonium bicarbonate and reduced at room temperature for 0.5 h. The DTT solution was removed and the sample alkylated in 30 µL 50 mM iodoacetamide in 0.1 M ammonium bicarbonate at room temperature for 0.5 h. The reagent was removed and the gel pieces dehydrated in 100 µL acetonitrile. The acetonitrile was removed and the gel pieces rehydrated in 100 µL 0.1 M ammonium bicarbonate. The pieces were dehydrated in 100 µL acetonitrile, the acetonitrile removed and the pieces completely dried by vacuum centrifugation. The gel pieces were rehydrated in 20 ng/µL (Trypsin, Chymotrypsin, and Thermolysin separately) in 50 mM ammonium bicarbonate on ice for 30 min. Any excess enzyme solution was removed and 20 µL 50 mM ammonium bicarbonate added. The sample was digested overnight at 37 $^{o}$C and the peptides formed extracted from the polyacrylamide in a 100 µL aliquot of 50% acetonitrile/5% formic acid. This extract was evaporated to 15 µL for MS analysis.

The LC-MS system consisted of a ThermoFisher Orbitrap Exploris 480 mass spectrometer system with an Easy Spray ion source connected to a Thermo 75 µm x 15 cm C18 Easy Spray column (through pre-column). 5 µL of the extract was injected and the peptides eluted from the column by an acetonitrile/0.1 M acetic acid gradient at a flow rate of 0.3 µL/min over 1.0 hours. The nanospray ion source was operated at 1.9 kV. The digest was analyzed using the rapid switching capability of the instrument acquiring a full scan mass spectrum to determine peptide molecular weights followed by product ion spectra (Top10 - 10 HCD) to determine amino acid sequence in sequential scans. This mode of analysis produces approximately 25000 MS/MS spectra of ions ranging in abundance over several orders of magnitude. Not all MS/MS spectra are derived from peptides. The data were analyzed by database searching using the Sequest search algorithm against investigator's provided sequence.

A

ELAALQGSEpT(230)PGKGGGSSSSSER

Peptide Summary

Sequence: ELAALQGSETPGKGGGSSSSSER, T10-Phospho (79.96633 Da)
Charge: +3, Monoisotopic m/z: 758.00769 Da (-0.12 mmu/-0.16 ppm), MH+: 2272.00852 Da, RT: 26.2921 min,
Identified with: Sequest HT (v1.17); XCorr:5.82, q-Value:7.2e-5, PEP:5.4e-8, PhosphoRS: Best Site Probabilities:T10(Phospho): 100,
Fragment match tolerance used for search: 0.02 Da
Fragments used for search: y-$H_2O$; y-$NH_3$; b; b-$H_2O$; b-$NH_3$; y

Fragment Matches

Value Type: Theo. Mass [Da]

Ion Series | Modification Losses | Neutral Losses | Multiple Neutral Losses | Precursor Ions | Internal Fragments

| #1 | $b^+$ | $b^{2+}$ | $b^{3+}$ | Seq. | $y^+$ | $y^{2+}$ | $y^{3+}$ | #2 |
|---|---|---|---|---|---|---|---|---|
| 1 | 130.04987 | 65.52857 | 44.02147 | E | | | | 23 |
| 2 | 243.13393 | 122.07061 | 81.71616 | L | 2142.96630 | 1071.98679 | 714.99362 | 22 |
| 3 | 314.17105 | 157.58916 | 105.39520 | A | 2029.88223 | 1015.44475 | 677.29893 | 21 |
| 4 | 385.20816 | 193.10772 | 129.07424 | A | 1958.84512 | 979.92620 | 653.61989 | 20 |
| 5 | 498.29223 | 249.64975 | 166.76893 | L | 1887.80800 | 944.40764 | 629.94085 | 19 |
| 6 | 626.35080 | 313.67904 | 209.45512 | Q | 1774.72394 | 887.86561 | 592.24616 | 18 |
| 7 | 683.37227 | 342.18977 | 228.46227 | G | 1646.66536 | 823.83632 | 549.55997 | 17 |
| 8 | 770.40429 | 385.70579 | 257.47295 | S | 1589.64390 | 795.32559 | 530.55282 | 16 |
| 9 | 899.44689 | 450.22708 | 300.48715 | E | 1502.61187 | 751.80957 | 501.54214 | 15 |
| 10 | 1080.46090 | 540.73409 | 360.82515 | T-Phospho | 1373.56928 | 687.28828 | 458.52794 | 14 |
| 11 | 1177.51366 | 589.26047 | 393.17607 | P | 1192.55527 | 596.78127 | 398.18994 | 13 |
| 12 | 1234.53512 | 617.77120 | 412.18323 | G | 1095.50250 | 548.25489 | 365.83902 | 12 |
| 13 | 1362.63009 | 681.81868 | 454.88155 | K | 1038.48104 | 519.74416 | 346.83186 | 11 |
| 14 | 1419.65155 | 710.32941 | 473.88870 | G | 910.38608 | 455.69668 | 304.13354 | 10 |
| 15 | 1476.67301 | 738.84015 | 492.89586 | G | 853.36461 | 427.18595 | 285.12639 | 9 |
| 16 | 1533.69448 | 767.35088 | 511.90301 | G | 796.34315 | 398.67521 | 266.11923 | 8 |
| 17 | 1620.72651 | 810.86689 | 540.91369 | S | 739.32169 | 370.16448 | 247.11208 | 7 |
| 18 | 1707.75854 | 854.38291 | 569.92436 | S | 652.28966 | 326.64847 | 218.10140 | 6 |
| 19 | 1794.79056 | 897.89892 | 598.93504 | S | 565.25763 | 283.13245 | 189.09073 | 5 |

★ site determining ions

Fragment Spectrum

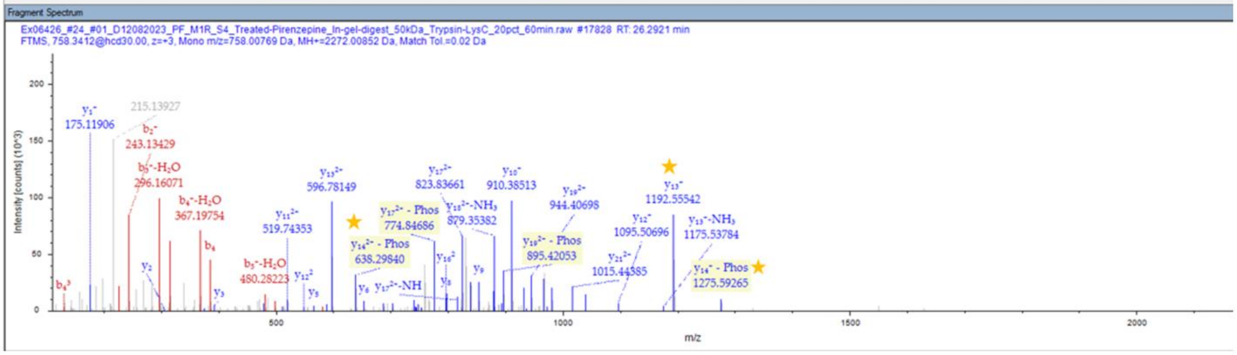


B

SQPGAEGpS(251)PETPPGR

Peptide Summary

Sequence: SQPGAEGSPETPPGR, S8-Phospho (79.96633 Da)
Charge: +2, Monoisotopic m/z: 773.83210 Da (+1.8 mmu/+2.32 ppm), MH+: 1546.65693 Da, RT: 20.8841 min,
Identified with: Sequest HT (v1.17); XCorr:3.80, q-Value:7.2e-5, PEP:6.2e-6, PhosphoRS: Best Site Probabilities:S8(Phospho): 99.57,
Fragment match tolerance used for search: 0.02 Da
Fragments used for search: y-$H_2O$; y-$NH_3$; b; b-$H_2O$; b-$NH_3$; y

Fragment Matches

Value Type: Theo. Mass [Da]

Ion Series | Modification Losses | Neutral Losses | Multiple Neutral Losses | Precursor Ions | Internal Fragments

| #1 | $b^+$ | $b^{2+}$ | Seq. | $y^+$ | $y^{2+}$ | #2 |
|---|---|---|---|---|---|---|
| 1 | 88.03930 | 44.52329 | S | | | 15 |
| 2 | 216.09788 | 108.55258 | Q | 1459.62131 | 730.31430 | 14 |
| 3 | 313.15065 | 157.07896 | P | 1331.56274 | 666.28501 | 13 |
| 4 | 370.17211 | 185.58969 | G | 1234.50997 | 617.75862 | 12 |
| 5 | 441.20922 | 221.10825 | A | 1177.48851 | 589.24789 | 11 |
| 6 | 570.25182 | 285.62955 | E | 1106.45139 | 553.72934 | 10 |
| 7 | 627.27328 | 314.14028 | G | 977.40880 | 489.20804 | 9 |
| 8 | 794.27164 | 397.63946 | S-Phospho | 920.38734 | 460.69731 | 8 |
| 9 | 891.32440 | 446.16584 | P | 753.38898 | 377.19813 | 7 |
| 10 | 1020.36700 | 510.68714 | E | 656.33621 | 328.67175 | 6 |
| 11 | 1121.41467 | 561.21098 | T | 527.29362 | 264.15045 | 5 |
| 12 | 1218.46744 | 609.73736 | P | 426.24594 | 213.62661 | 4 |
| 13 | 1315.52020 | 658.26374 | P | 329.19318 | 165.10023 | 3 |
| 14 | 1372.54167 | 686.77447 | G | 232.14042 | 116.57385 | 2 |
| 15 | | | R | 175.11895 | 88.06311 | 1 |

Fragment Spectrum

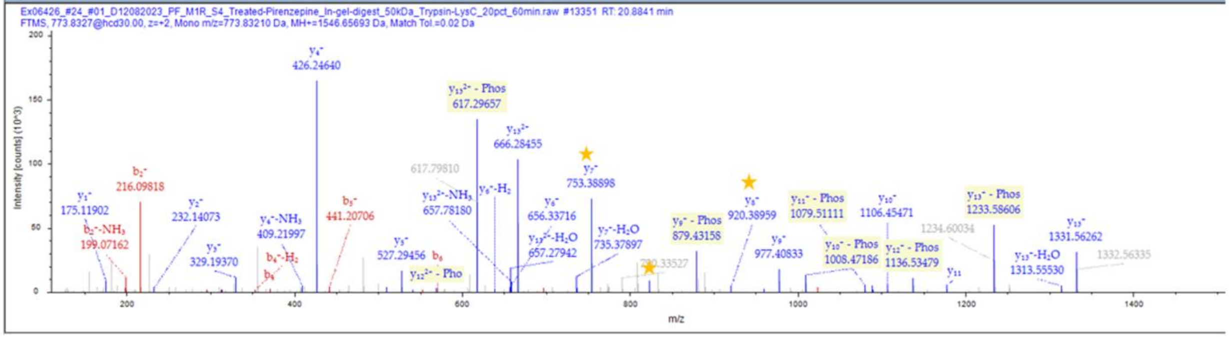

C

SQPGAEGSPEpT(254)PPGR

Peptide Summary

Sequence: SQPGAEGSPETPPGR, T11-Phospho (79.96633 Da)
Charge: +2, Monoisotopic m/z: 773.83000 Da (-0.31 mmu/-0.4 ppm), MH+: 1546.65272 Da, RT: 21.4967 min,
Identified with: Sequest HT (v1.17); XCorr:4.23, q-Value:7.2e-5, PEP:2.8e-6, PhosphoRS: Best Site Probabilities:T11(Phospho): 100,
Fragment match tolerance used for search: 0.02 Da
Fragments used for search: y-H₂O; y-NH₃; b; b-H₂O; b-NH₃; y

Fragment Matches

Value Type: Theo. Mass [Da]

Ion Series | Modification Losses | Neutral Losses | Multiple Neutral Losses | Precursor Ions | Internal Fragments

| #1 | $b^+$ | $b^{2+}$ | Seq. | $y^+$ | $y^{2+}$ | #2 |
|---|---|---|---|---|---|---|
| 1 | 88.03930 | 44.52329 | S | | | 15 |
| 2 | 216.09788 | 108.55258 | Q | 1459.62131 | 730.31430 | 14 |
| 3 | 313.15065 | 157.07896 | P | 1331.56274 | 666.28501 | 13 |
| 4 | 370.17211 | 185.58969 | G | 1234.50997 | 617.75862 | 12 |
| 5 | 441.20922 | 221.10825 | A | 1177.48851 | 589.24789 | 11 |
| 6 | 570.25182 | 285.62955 | E | 1106.45139 | 553.72934 | 10 |
| 7 | 627.27328 | 314.14028 | G | 977.40880 | 489.20804 | 9 |
| 8 | 714.30531 | 357.65629 | S | 920.38734 | 460.69731 | 8 |
| 9 | 811.35807 | 406.18267 | P | 833.35531 | 417.18129 | 7 |
| 10 | 940.40067 | 470.70397 | E | 736.30255 | 368.65491 | 6 |
| 11 | 1121.41467 | 561.21098 | T-Phospho | 607.25995 | 304.13361 | 5 |
| 12 | 1218.46744 | 609.73736 | P | 426.24594 | 213.62661 | 4 |
| 13 | 1315.52020 | 658.26374 | P | 329.19318 | 165.10023 | 3 |
| 14 | 1372.54167 | 686.77447 | G | 232.14042 | 116.57385 | 2 |
| 15 | | | R | 175.11895 | 88.06311 | 1 |

Fragment Spectrum

Ex06426_#24_#01_D12082023_PF_M1R_S4_Treated-Pirenzepine_In-gel-digest_50kDa_Trypsin-LysC_20pct_60min.raw #13791 RT: 21.4967 min
FTMS, 773.8306@hcd30.00, z=+2, Mono m/z=773.83000 Da, MH+=1546.65272 Da, Match Tol.=0.02 Da

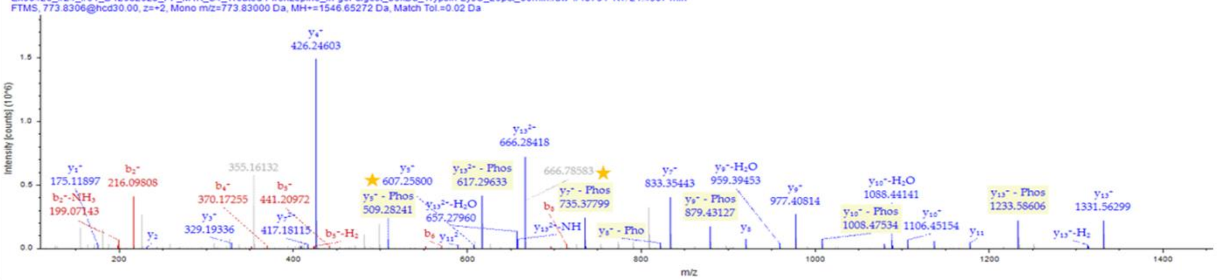


D

MPMVDPEAQAPTKQPPRpS(321)SPNTVK

Peptide Summary

Sequence: MPMVDPEAQAPTKQPPRSSPNTVK, S18-Phospho (79.96633 Da)
Charge: +4, Monoisotopic m/z: 672.32661 Da (+2.95 mmu/+4.38 ppm), MH+: 2686.28462 Da, RT: 28.6018 min,
Identified with: Sequest HT (v1.17); XCorr:4.24, q-Value:4.5e-5, PEP:2.1e-5, PhosphoRS: Best Site Probabilities:S18(Phospho): 98.2,
Fragment match tolerance used for search: 0.02 Da
Fragments used for search: y-H₂O; y-NH₃; b; b-H₂O; b-NH₃; y

Fragment Matches

Value Type: Theo. Mass [Da]

Ion Series | Modification Losses | Neutral Losses | Multiple Neutral Losses | Precursor Ions | Internal Fragments

| #1 | $b^+$ | $b^{2+}$ | $b^{3+}$ | Immonium | Seq. | $y^+$ | $y^{2+}$ | $y^{3+}$ | #2 |
|---|---|---|---|---|---|---|---|---|---|
| 1 | 132.04776 | 66.52752 | 44.68744 | 104.05285 | M | | | | 24 |
| 2 | 229.10053 | 115.05390 | 77.03836 | 70.06513 | P | 2555.23236 | 1278.11982 | 852.41564 | 23 |
| 3 | 360.14101 | 180.57414 | 120.71852 | 104.05285 | M | 2458.17960 | 1229.59344 | 820.06472 | 22 |
| 4 | 459.20942 | 230.10835 | 153.74133 | 72.08078 | V | 2327.13912 | 1164.07320 | 776.38456 | 21 |
| 5 | 574.23637 | 287.62182 | 192.08364 | 88.03930 | D | 2228.07070 | 1114.53899 | 743.36175 | 20 |
| 6 | 671.28913 | 336.14820 | 224.43456 | 70.06513 | P | 2113.04376 | 1057.02552 | 705.01944 | 19 |
| 7 | 800.33172 | 400.66950 | 267.44876 | 102.05496 | E | 2015.99099 | 1008.49914 | 672.66852 | 18 |
| 8 | 871.36884 | 436.18806 | 291.12780 | 44.04948 | A | 1886.94840 | 943.97784 | 629.65432 | 17 |
| 9 | 999.42741 | 500.21735 | 333.81399 | 101.07094 | Q | 1815.91129 | 908.45928 | 605.97528 | 16 |
| 10 | 1070.46453 | 535.73590 | 357.49303 | 44.04948 | A | 1687.85271 | 844.42999 | 563.28909 | 15 |
| 11 | 1167.51729 | 584.26228 | 389.84395 | 70.06513 | P | 1616.81560 | 808.91144 | 539.61005 | 14 |
| 12 | 1268.56497 | 634.78612 | 423.52651 | 74.06004 | T | 1519.76283 | 760.38505 | 507.25913 | 13 |
| 13 | 1396.65993 | 698.83360 | 466.22483 | 101.10732 | K | 1418.71515 | 709.86122 | 473.57657 | 12 |
| 14 | 1524.71851 | 762.86289 | 508.91102 | 101.07094 | Q | 1290.62019 | 645.81373 | 430.87825 | 11 |
| 15 | 1621.77127 | 811.38928 | 541.26194 | 70.06513 | P | 1162.56161 | 581.78445 | 388.19206 | 10 |
| 16 | 1718.82404 | 859.91566 | 573.61286 | 70.06513 | P | 1065.50885 | 533.25806 | 355.84113 | 9 |
| 17 | 1874.92515 | 937.96621 | 625.64657 | 129.11347 | R | 968.45609 | 484.73168 | 323.49021 | 8 |
| 18 | 2041.92351 | 1021.46539 | 681.31269 | 140.01072 | S-Phospho | 812.35498 | 406.68113 | 271.45651 | 7 |
| 19 | 2128.95554 | 1064.98141 | 710.32336 | 60.04439 | S | 645.35662 | 323.18195 | 215.79039 | 6 |
| 20 | 2226.00830 | 1113.50779 | 742.67428 | 70.06513 | P | 558.32459 | 279.66593 | 186.77971 | 5 |
| 21 | 2340.05123 | 1170.52925 | 780.68859 | 87.05529 | N | 461.27182 | 231.13955 | 154.42879 | 4 |
| 22 | 2441.09891 | 1221.05309 | 814.37115 | 74.06004 | T | 347.22890 | 174.11809 | 116.41448 | 3 |
| 23 | 2540.16732 | 1270.58730 | 847.39396 | 72.08078 | V | 246.18122 | 123.59425 | 82.73192 | 2 |
| 24 | | | | 101.10732 | K | 147.11280 | 74.06004 | 49.70912 | 1 |

Fragment Spectrum

Ex05075_#24_#01_D12082022_PF_M1R_MT7_S-trap_LysC_45K_150ng_60min.raw #13294 RT: 28.6018 min
FTMS, 672.5775@hcd30.00, z=+4, Mono m/z=672.32661 Da, MH+=2686.28462 Da, Match Tol.=0.02 Da

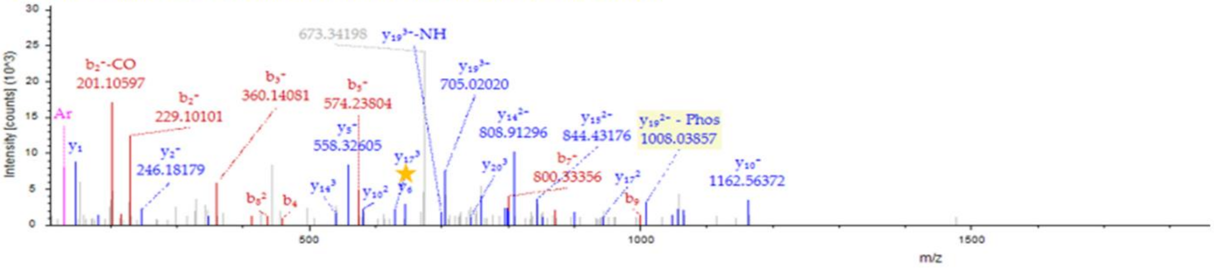

E

pT(354)FSLVKEK

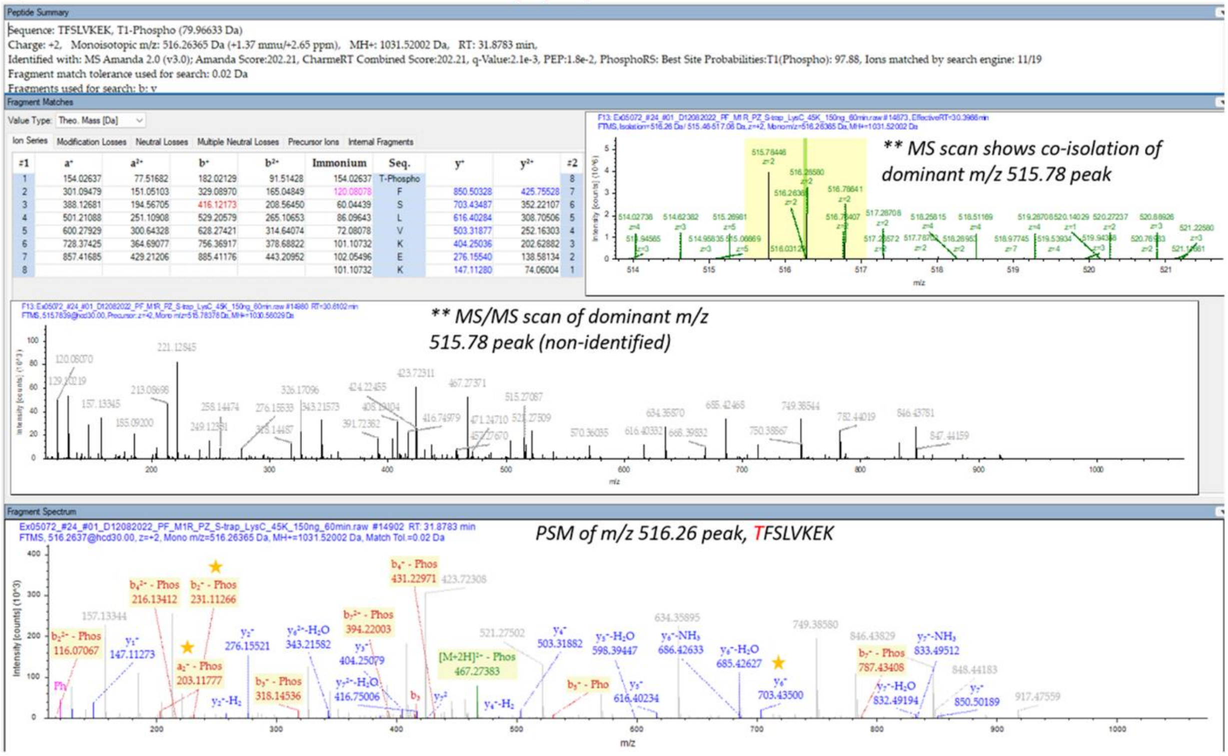


| #1 | a⁺ | a²⁺ | b⁺ | b²⁺ | Immonium | Seq. | y⁺ | y²⁺ | #2 |
|---|---|---|---|---|---|---|---|---|---|
| 1 | 154.02637 | 77.51682 | 182.02129 | 91.51428 | 154.02637 | T-Phospho | | | 8 |
| 2 | 301.09479 | 151.05103 | 329.08970 | 165.04849 | 120.08078 | F | 850.50328 | 425.75528 | 7 |
| 3 | 388.12681 | 194.56705 | 416.12173 | 208.56450 | 60.04439 | S | 703.43487 | 352.22107 | 6 |
| 4 | 501.21088 | 251.10908 | 529.20579 | 265.10653 | 86.09643 | L | 616.40284 | 308.70506 | 5 |
| 5 | 600.27929 | 300.64328 | 628.27421 | 314.64074 | 72.08078 | V | 503.31877 | 252.16303 | 4 |
| 6 | 728.37425 | 364.69077 | 756.36917 | 378.68822 | 101.10732 | K | 404.25036 | 202.62882 | 3 |
| 7 | 857.41685 | 429.21206 | 885.41176 | 443.20952 | 102.05496 | E | 276.15540 | 138.58134 | 2 |
| 8 | | | | | 101.10732 | K | 147.11280 | 74.06004 | 1 |

F

TFpS(356)LVK

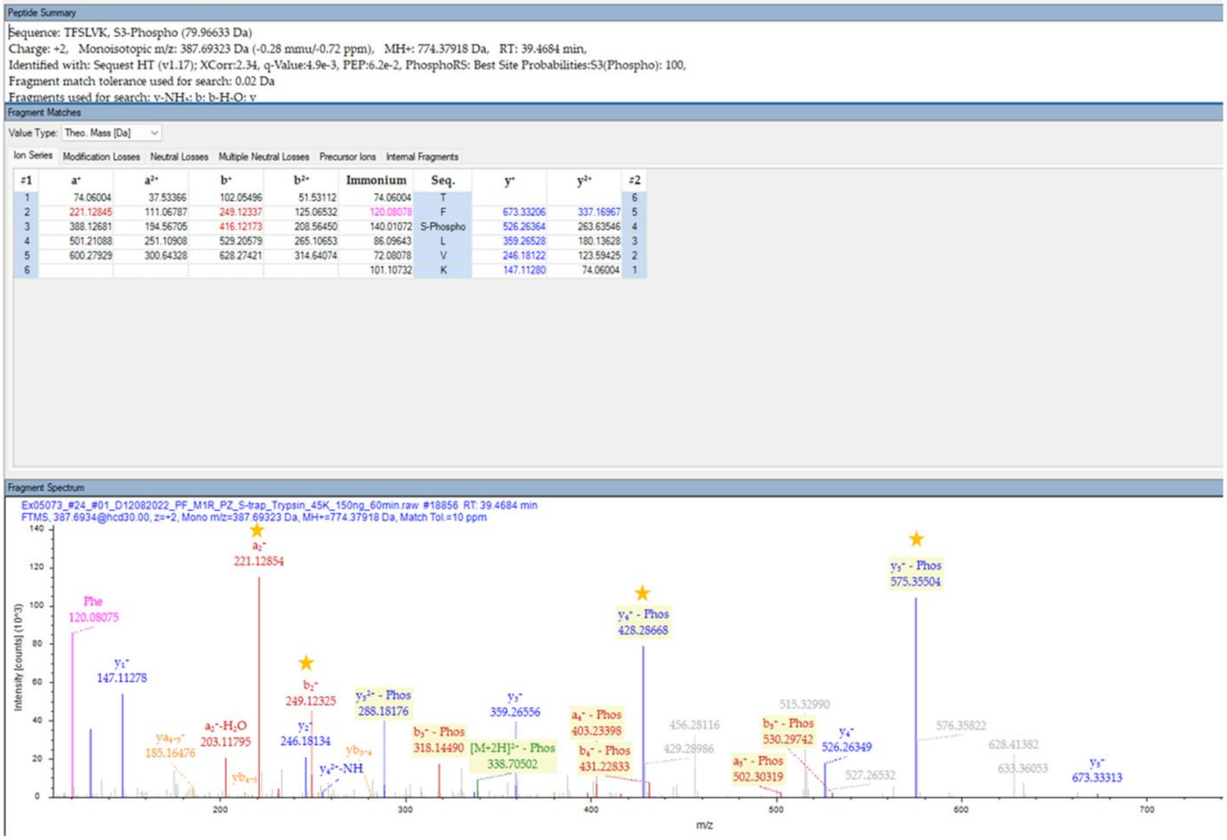


| #1 | a⁺ | a²⁺ | b⁺ | b²⁺ | Immonium | Seq. | y⁺ | y²⁺ | #2 |
|---|---|---|---|---|---|---|---|---|---|
| 1 | 74.06004 | 37.53366 | 102.05496 | 51.53112 | 74.06004 | T | | | 6 |
| 2 | 221.12845 | 111.06787 | 249.12337 | 125.06532 | 120.08078 | F | 673.33206 | 337.16967 | 5 |
| 3 | 388.12681 | 194.56705 | 416.12173 | 208.56450 | 140.01072 | S-Phospho | 526.26364 | 263.63546 | 4 |
| 4 | 501.21088 | 251.10908 | 529.20579 | 265.10653 | 86.09643 | L | 359.26528 | 180.13628 | 3 |
| 5 | 600.27929 | 300.64328 | 628.27421 | 314.64074 | 72.08078 | V | 246.18122 | 123.59425 | 2 |
| 6 | | | | | 101.10732 | K | 147.11280 | 74.06004 | 1 |

**Fig. S3. LC-MS/MS analysis identifies sites of $hM_1R$ phosphorylation in HEK293 cells following pirenzepine treatment.** LC-MS/MS analysis performed by the Manitoba Centre for Proteomics and Systems Biology (MCPSB) **(A**) MS2 spectra of identified $hM_1R$ phosphorylation at T230, **(B)** at S251, **(C)** at T254, **(D)** at S321, **(E)** at T354, and **(F)** at S356 in ICL3.

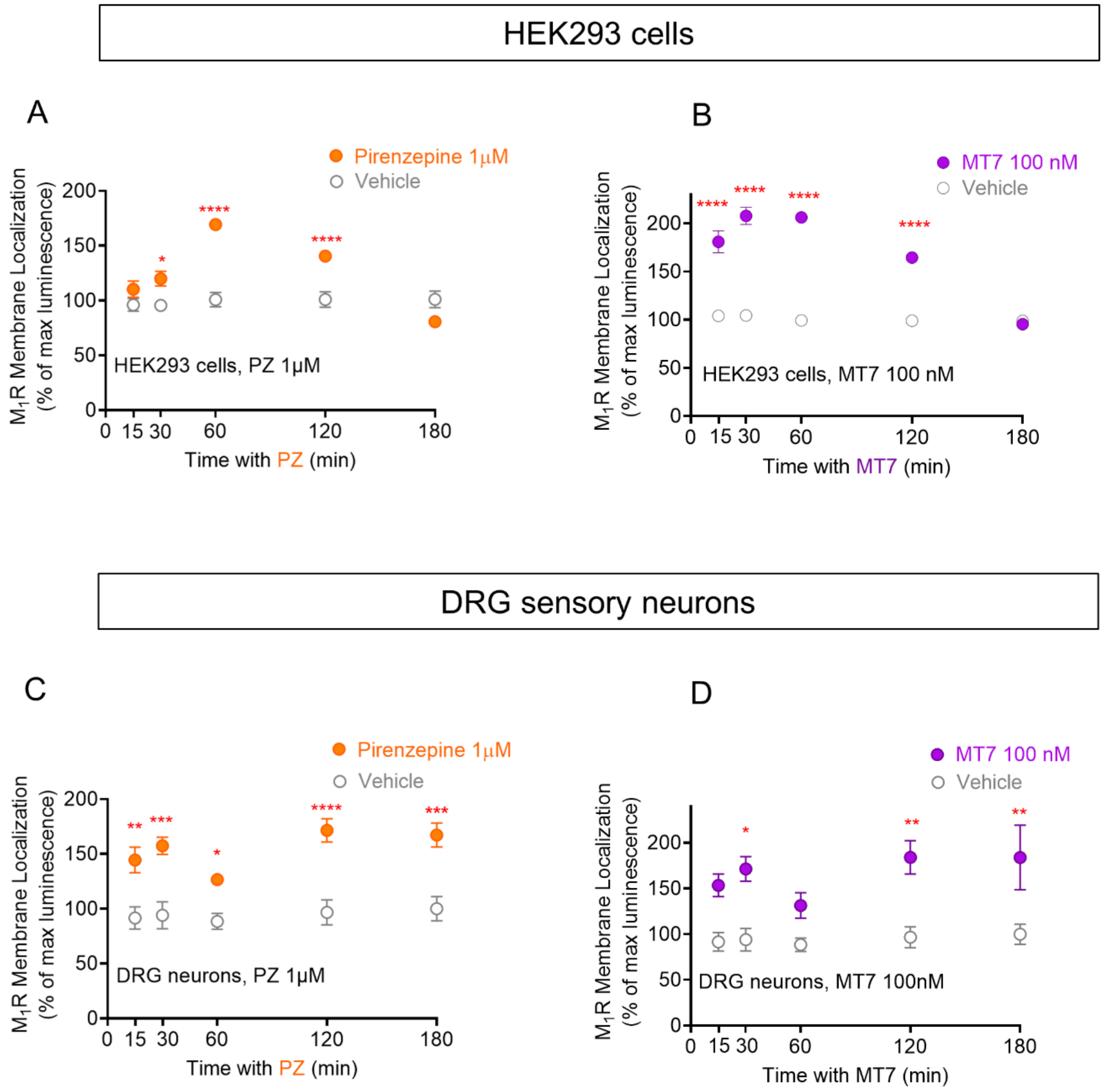


**Fig. S4. Time course of PZ and MT7 induced accumulation of $M_1$R at the cell surface.** Time-response curve for receptor internalization performed for **(A)** pirenzepine (PZ, 1 µM, HEK293 cells), **(B)** MT7 (100 nM, HEK293 cells), **(C)** pirenzepine (PZ, 1 µM, DRG sensory neurons), and **(D)** MT7 (100 nM, DRG sensory neurons). *t*-test was used to compare the values of drug-treated groups with their vehicle-treated groups at each time-point. Data show mean ± SEM; ****P<0.0001, ***P<0.001, **P<0.01 and *P<0.05, n=4.

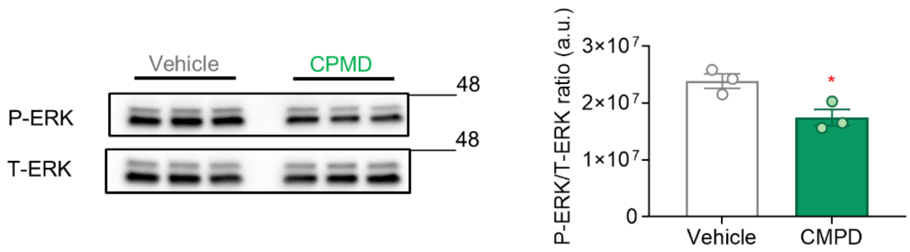


**Fig. S5. CPMD101 treatment in non-transfected HEK293 cells.** HEK293 cells received vehicle/CMPD101 (30 µM, 10 min) and immunoblotting for T-ERK and P-ERK proteins was conducted. Data represent P-ERK/T-ERK ratio (a.u.). Data show mean ± SEM; *t*-test, *P<0.05, n=3.

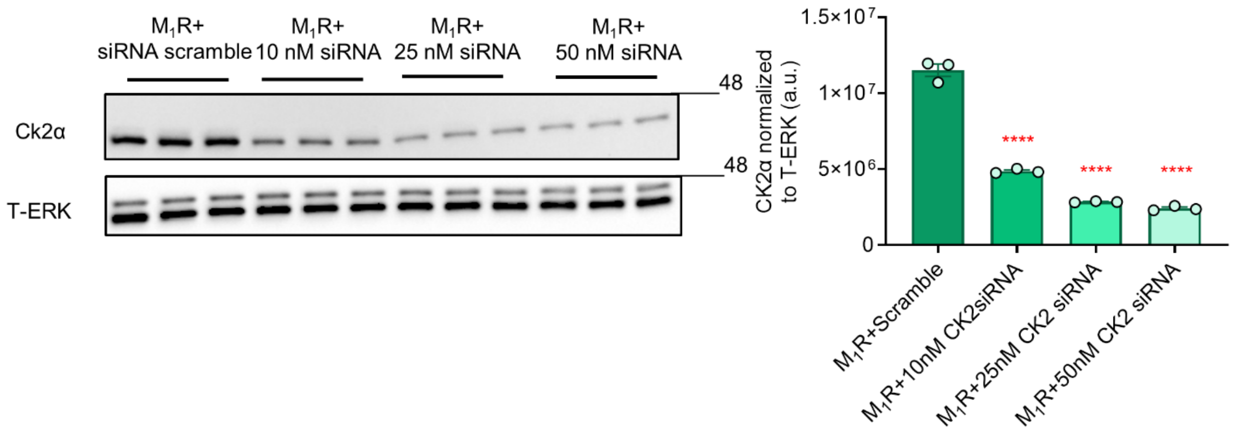


**Fig. S6. CK2α knock-down in HEK293 cells.** HEK293 cells received scramble siRNA (50 nM) or target pool siRNA for CK2α at different concentrations (10, 25, and 50 nM) for 24h, and then were transfected with $hM_1R$ plasmid (100 ng/well) for another 24h in 6-well plates. Immunoblotting for CK2α and T-ERK proteins was conducted. Data represent CK2α values normalized to T-ERK (a.u.). Data show mean ± SEM (1-way ANOVA with Dunnett's post-hoc test). Comparison to siRNA scramble; ****P<0.0001, n=3.

### Tables S1 and S2

| Annotated Sequence | m/z [Da] | MH+ [Da] | Site |
|---|---|---|---|
| SQPGAEGSPEtPPGR | 773.8308 | 1546.6542 | T254 |
| SQPGAEGsPETPPGR | 773.8309 | 1546.6545 | S251 |

**Table. S1. Table of identified phosphorylated peptides on $hM_1R$.** Phosphorylated residues were determined by label-free LC-MS/MS on trypsin-digested and phosphorylation-enriched in HEK293 cells following overexpression of $hM_1R$ (24h) and treatment with pirenzepine (1 µM, 30 min). Results from Biomolecular Analysis Facility at University of Virginia Health System.

| Annotated Sequence | m/z [Da] | MH+ [Da] | Site |
|---|---|---|---|
| ELAALQGSEtPGKGGGSSSSSER | 758.0076 | 2272.0085 | T230 |
| SQPGAEGsPETPPGR | 773.8321 | 1546.6569 | S251 |
| SQPGAEGSPEtPPGR | 773.8300 | 1546.6527 | T254 |
| MPMVDPEAQAPTKQPPRsSPNTVK | 672.3266 | 2686.2846 | S321 |
| tFSLVKEK | 516.2636 | 1031.520 | T354 |
| TFsLVK | 387.6932 | 774.3791 | S356 |

**Table. S2. Table of identified phosphorylated peptides on $hM_1R$.** Phosphorylated residues were determined by label-free LC-MS/MS on trypsin-digested and phosphorylation-enriched in HEK293 cells following overexpression of $hM_1R$ (24h) and treatment with pirenzepine (1 µM, 30 min). Results from Manitoba Centre for Proteomics and Systems Biology (MCPSB).
